# An algorithm for identifying task-specific brain subnetworks using the visuomotor system as an example

**DOI:** 10.1101/2024.05.13.593972

**Authors:** Ryan Ellison, Mona Matar, Suleyman A. Gokoglu, Raj K. Prabhu, Scott L. Hooper

**Affiliations:** Universities Space Research Association, Cleveland, OH, USA; Department of Biological Sciences, Ohio University, Athens, OH, USA; NASA Glenn Research Center, Cleveland, OH, USA

**Keywords:** visuomotor, module, subnetwork, connectome, macroscopic model, clustering

## Abstract

We describe an algorithm that identifies a subnetwork of brain regions involved in producing a task-specific behavior, here visuomotor behavior, from an anatomically defined primate brain connectome. The algorithm first finds the brain regions connected to an output region (here, primary motor cortex, M1) by one connection. It then identifies all regions, termed “layer 2” regions, connected to these “layer 1” regions by one connection. This process continues until the layer containing the input region (here, primary visual cortex, V1) is reached. The algorithm then finds, subject to a user-set maximum step number, all paths linking the input and output regions. The brain regions in these paths constitute the initial subnetwork identification that performs the task. Regions known not to be task-involved (for example, regions in the ventral stream of visual information vs. the dorsal stream, which helps generate visuomotor behavior) are then removed. Structural subnetwork analysis showed that the intraparietal sulcus of the parietal cortex (PCIP) was most, and the secondary visual (V2) and superior parietal (PCS) cortices second-most, central to local network activity. Changing PCIP, V2 and PCS activity was thus most likely to alter activity of the entire subnetwork. Model sufficiency was tested by instantiating each brain region’s inherent activity with multiple versions of a simple two-dimensional (2D) model that can produce oscillatory activity and synaptically interconnecting the regions to produce a macroscopic visuomotor model. The model reproduced the experimental local field potential (LFP) activity of the brain regions identified as part of the visuomotor subnetwork.

## 1 Introduction

An unresolved question in neurobiology is how to abstract from global maps of long-distance connectivity among brain regions—connectomes connectomes—the specific set of brain regions active during expression of a behavior (e.g., reach-to-grasp-like movements). In some large networks, structural modularity methods from graph theory such as partitioning (Blondel et al., 2008; Clauset et al., 2004; Duch and Arenas, 2005; Girvan and Newman, 2002; Guimerà and Nunes Amaral, 2005; Newman, 2004; Newman and Girvan, 2004) and distance-based methods (Hastie et al., 2009) can find clusters of densely interconnected nodes with sparse inter-cluster connectivity (Arenas et al., 2006; Sporns and Betzel, 2016; Wildie and Shanahan, 2012). These subnetworks are called communities or modules and can produce complex activity including metastability and synchronizability (Fortunato, 2010). Applying these methods to whole-brain connectomes is difficult because their efficacy depends on module conceptualization and characteristics of the connectivity data (Sporns and Betzel, 2016), and the 76 brain regions in the primate connectome (Leon et al., 2013; Leon et al., 2015) is markedly less than that needed to identify meaningful modules (Fortunato and Barthélemy, 2007).

Moreover, modules identified in this manner are not necessarily those involved in producing any particular behavior. An approach that combined connectome and experimental data on brain region function to identify task-specific functional brain region subnetworks would therefore be useful. For many behaviors, the task is to produce a motor output in response to a sensory input, which immediately identifies an output (primary motor cortex, M1) and input (the initial cortical area involved in the relevant sensory modality) brain region. The putative functional brain region subnetwork would be the set of regions most interconnected between the input and output regions. We are unaware of this approach being used before and present here an algorithm to identify such task-dependent functional subnetworks.

Once a putative functional brain region subnetwork has been thus identified, the inherent activity of each brain region in the subnetwork must be specified. EEG recordings of entire brain activity show multiple frequency peaks divided into delta (0.5-4 Hz), theta (4-8 Hz), alpha (8-12 Hz), beta (12-35 Hz), and gamma (>35 Hz) ranges (Abhang et al., 2016). Local field potential (LFP) extracellular recordings are the summed activity of an individual brain region’s many neurons acting in synchrony. Summing of these brain region LFPs as a function of distance between the brain region and the EEG electrode gives rise to EEG recordings (Destexhe and Bedard, 2013). LFP recordings show that, when active, individual brain regions oscillate at characteristic frequencies.

This synchronized, single frequency oscillation allows modeling each region as a unit oscillator (Jansen and Rit, 1995; Leon et al., 2015; Stefanescu and Jirsa, 2008; 2011; Wilson and Cowan, 1972; 1973; Wong and Wang, 2006). An often-used approach, and that used here, is to model individual brain regions with a 2D system (i.e., a system with two ordinary differential equations) generalized from simplified models of individual neurons (Fitzhugh, 1961; Hindmarsh and Rose, 1984; Izhikevich, 2003; Nagumo et al., 1962; Rinzel, 1987; Van der Pol, 1926; Wilson, 1981) and neuronal populations (Jansen and Rit, 1995; Leon et al., 2015; Stefanescu and Jirsa, 2008; 2011; Wilson and Cowan, 1972; 1973; Wong and Wang, 2006). This approach successfully models resting state brain activity (Deco et al., 2017; Ghosh et al., 2008) and has led to the idea that biological neural networks at rest are near state transitions, a concept called self-organized criticality (Chialvo, 2010; Hesse and Gross, 2014). However, to our knowledge, it has not been applied to populate task-dependent brain region subnetworks.

Applying these ideas to the brain connectome using data from visuomotor behaviors such as reach-to-grasp (Berens et al., 2008; Frien et al., 1994; MacKay and Mendonca, 1995; Murthy and Fetz, 1992; Rougeul et al., 1979; Sanes and Donoghue, 1993; Stetson and Andersen, 2014; Vaidya et al., 2015; Weiss et al., 2018; Womelsdorf et al., 2010) identified a 14-region visuomotor subnetwork. Applying graph theory analysis techniques to the subnetwork identified the intraparietal sulcus of the parietal cortex (PCIP), the secondary visual cortex (V2), and the superior parietal cortex (PCS) as central regions of the subnetwork. Changes in PCIP, V2 and PCS activity are therefore most likely to alter the activity of the entire subnetwork. Multiple instantiations of unit oscillator models that reproduced each region’s oscillatory frequency when active were then introduced into the 14-region visuomotor subnetwork. When in the subnetwork, the “intrinsic” activities of the regions shifted but remained within the frequency ranges identified from experimental macroscopic recordings and continued to do so for all instantiations of each region’s intrinsic 2D model. Moreover, the brain region LFP activities were coordinated to produce stereotypical weakly-coupled phase relationships (1:1 and n:m coupling, as well as phasic drift) as is commonly observed in neural networks at the single cell level.

## 2 Materials and Methods

This article describes a new algorithm for identifying functional brain region subnetworks. Most of the explanation of the algorithm is therefore in the Results. We describe here only the connectome dataset used as input to the algorithm, the algorithm pseudocode, the graph theory techniques used to analyze the structural visuomotor subnetwork, and implementation details of the generic brain region model (see **Section 3.3 A two-dimensional model that produces a wide range of outputs**, **Section 3.4 Incorporating brain region models into the visuomotor functional subnetwork**, Figure 9, and Table 2 for more modeling details).

### 2.1 Primate connectome data

The connectivity dataset for brain region identity, spatial location, and synaptic connectivity strength and length was acquired from The Virtual Brain (TVB) simulation platform, which pre-packages a macroscopic connectome dataset of a primate brain (Leon et al., 2013; Leon et al., 2015). This dataset contains 76 cortical brain regions (Table 1), represented as nodes, and 1,560 long-range, interregional connections (e.g., corticocortical connections, commissural fibers, and association tracts) represented as edges. The nodes are simple location data (each brain region’s three-dimensional center of mass), as opposed to, for example, the vertex-based mesh used to model cortical surfaces. This prepackaged dataset is itself derived from a large number of sources (Aboitiz, 1992; Bakker et al., 2012; Bezgin et al., 2009; Corballis, 2017; Hugdahl, 2005; Kötter, 2004; Kötter and Wanke, 2005; Leon et al., 2013; Leon et al., 2015; Paxinos et al., 2000; Stephan, 2013; Toga et al., 2009).

**Table 1.** Abbreviations of 76 brain regions in primate connectome. Only 38 listed because each region is in both left and right hemispheres.

| Abbreviation | Region | Abbreviation | Region |
| --- | --- | --- | --- |
| A1 | Primary auditory cortex | PFCDM | Prefrontal cortex dorsomedial |
| A2 | Secondary auditory cortex | PFCM | Prefrontal cortex medial |
| AMYG | amygdala | PFCORB | Prefrontal cortex orbital |
| CCA | Cingulate cortex anterior | PFCPOL | Prefrontal cortex pole |
| CCP | Cingulate cortex posterior | PFCVL | Prefrontal cortex ventrolateral |
| CCR | Cingulate cortex retrosplenialis | PHC | Parahippocampal cortex |
| CCS | Cingulate cortex subgenualis | PMCDL | Premotor cortex dorsolateral |
| FEF | Frontal eye field | PMCM | Premotor cortex medial<br>(supplementary motor cortex) |
| G | Gustatory cortex | PMCVL | Premotor cortex ventrolateral |
| HC | Hippocampal cortex | S1 | Primary somatosensory cortex |
| IA | Insula anterior | S2 | Secondary somatosensory<br>cortex |
| IP | Insula posterior | TCC | Temporal cortex central |
| M1 | Primary motor cortex | TCI | Temporal cortex inferior |
| PCI | Parietal cortex inferior | TCPOL | Temporal cortex pole |
| PCIP | Parietal cortex intraparietal sulcus | TCS | Temporal cortex superior |
| PCM | Parietal cortex medial | TCV | Temporal cortex ventral |
| PCS | Parietal cortex superior | V1 | Primary visual cortex |
| PFCCL | Prefrontal cortex centrolateral | V2 | Secondary visual cortex |
| PFCDL | Prefrontal cortex dorsolateral | CC | Corpus callosum |

**Table 2.** Abbreviations and expanded names of 14 brain regions in visuomotor module and of four hand-pruned regions.

| Module | Pruned | Region |
| --- | --- | --- |
| CCA |  | Cingulate cortex anterior |
| CCP |  | Cingulate cortex posterior |
| IP |  | Insula posterior |
| M1 |  | Primary motor cortex |
| PCIP |  | Parietal cortex intraparietal sulcus |
| PCM |  | Parietal cortex medial |
| PCS |  | Parietal cortex superior |
| PMCDL |  | Premotor cortex dorsolateral |
| PMCM |  | Premotor cortex medial (supplementary motor cortex) |
| PMCVL |  | Premotor cortex ventrolateral |
| S1 |  | Primary somatosensory cortex |
| S2 |  | Secondary somatosensory cortex |
| V1 |  | Primary visual cortex |
| V2 |  | Secondary visual cortex |
|  | A | amygdala |
|  | PFCORB | Prefrontal cortex orbital |
|  | PFCPOL | Prefrontal cortex pole |
|  | TCC | Temporal cortex central |

Figure 1 summarizes the whole-brain, primate connectome. Figure 1A shows the locations of the three-dimensional centers of mass of the brain regions with a generic representation of the brain surface. Figure 1B is a tract-synaptic weight matrix (76 x 76) whose entries are the synaptic weight between each brain region pair (a synaptic weight of zero indicates no synaptic connection; note that regions can make recursive synapses onto themselves and thus the diagonals in each quadrant are not necessarily zero). In this matrix columns are the source regions and rows the target regions; that is, the entry in column region 1, row region 2 is the synaptic weight from region 1 to region 2. In general, this is not the same as the weight from region 2 to region 1 (the entry in column region 2, row region 1), meaning this matrix is asymmetric within a hemisphere (although it is symmetric across hemispheres). Intra-hemisphere connectivity (first and third quadrants) is dense but inter-hemisphere connectivity (second and fourth quadrants) is sparse because only relatively few brain regions connect to themselves in the contralateral hemisphere.

**Figure 1.**
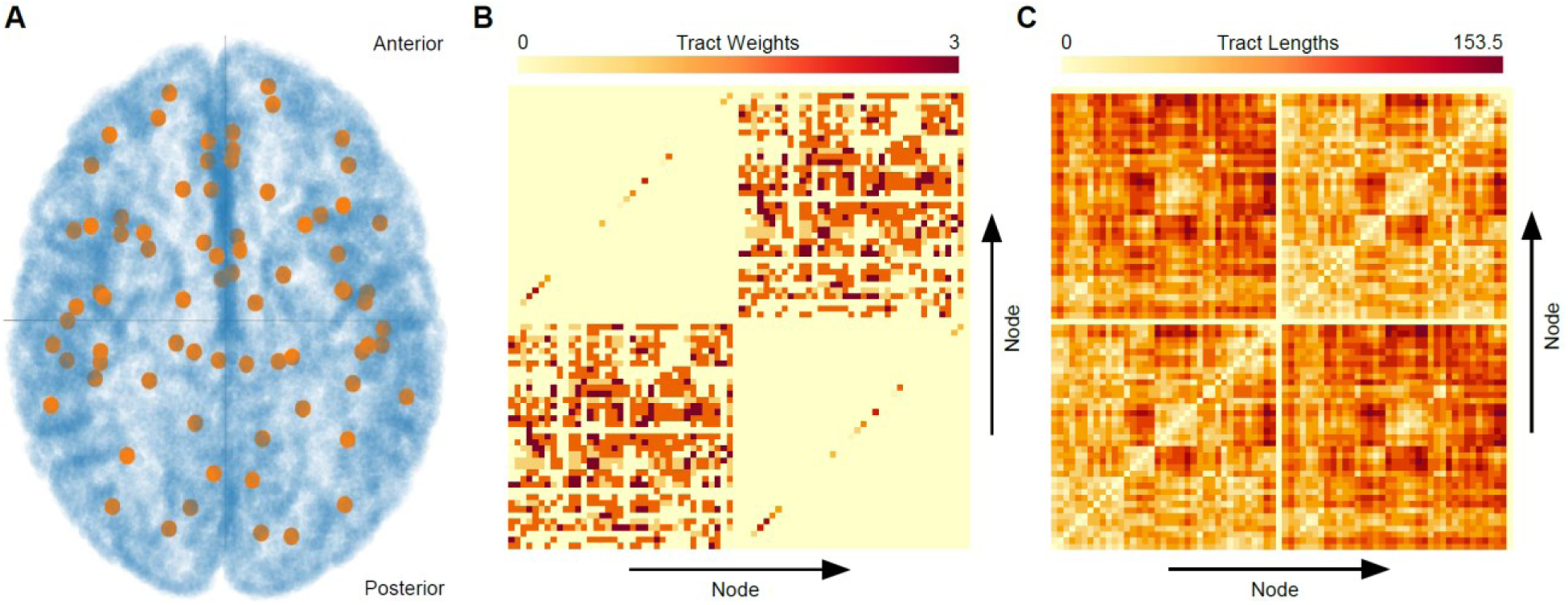
Primate connectome. **A.** A two-dimensional representation of the TVB data (76 nodes, orange filled circles) with cortical surface (blue) illustrating the diffuse spatiality and granularity of brain regions. 1,560 interregional connections not shown for clarity. Brain orientation: superior-to-inferior along axial plane, anterior-to-posterior top to bottom. **B.** The 76 x 76 tract-synaptic weight matrix with main diagonal from bottom left to top right. The upper right quadrant is all connections between regions in the right hemisphere, the lower left all connections between regions in the left hemisphere, the upper left all connections between left to right hemispheres, and the lower right all connections between right to left hemispheres. The inter-hemisphere connections are only those between the same brain region on the two sides and hence has non-zero values on the matrix diagonal. This matrix represents a weighted, undirected graph. **C.** The 76 x 76 tract-connection length matrix. Quadrants have same descriptions as in **B**. This asymmetric matrix represents a weighted, directed graph.

Figure 1C is a tract-length matrix (76 x 76) whose entries are the physical distances between each brain region pair (in each hemisphere each region therefore has a zero distance from itself; note that the diagonals in the upper right and lower left quadrants are all zero). Since these distances are the same regardless of source (column) and target (row) region, this matrix is symmetric within a hemisphere (because of differences of brain region localization in the two hemispheres, it is asymmetric across hemispheres).

### 2.2 Algorithm pseudocode

We describe the logic of finding the visuomotor subnetwork with a fully worked through example in the Results (see Figure 6). Here we present the pseudocode of particularly important parts of the algorithm. The process began with algorithm initialization and finding all regions connected to the output region, M1 (“layer 1”; Figures 2; see also **Section 3.1.2 Identifying (unpacking) all layers of regions between output and input region**). The algorithm then finds all regions connected to these regions which, for the visuomotor functional subnetwork, contained the input region, V1 (“layer 2”; Figure 3). We call this procedure the “unpacking” function. The algorithm then finds all pathways from the input region that connect to the output region (the “rebuild” function), subject to a user-set constraint on how long such pathways can be (i.e., how many steps the algorithm can take; Figure 4; see also **Section 3.1.3 Identifying (rebuilding) a pathway between the input and output region (node) subject to the connection length restriction**).

**Figure 2.**
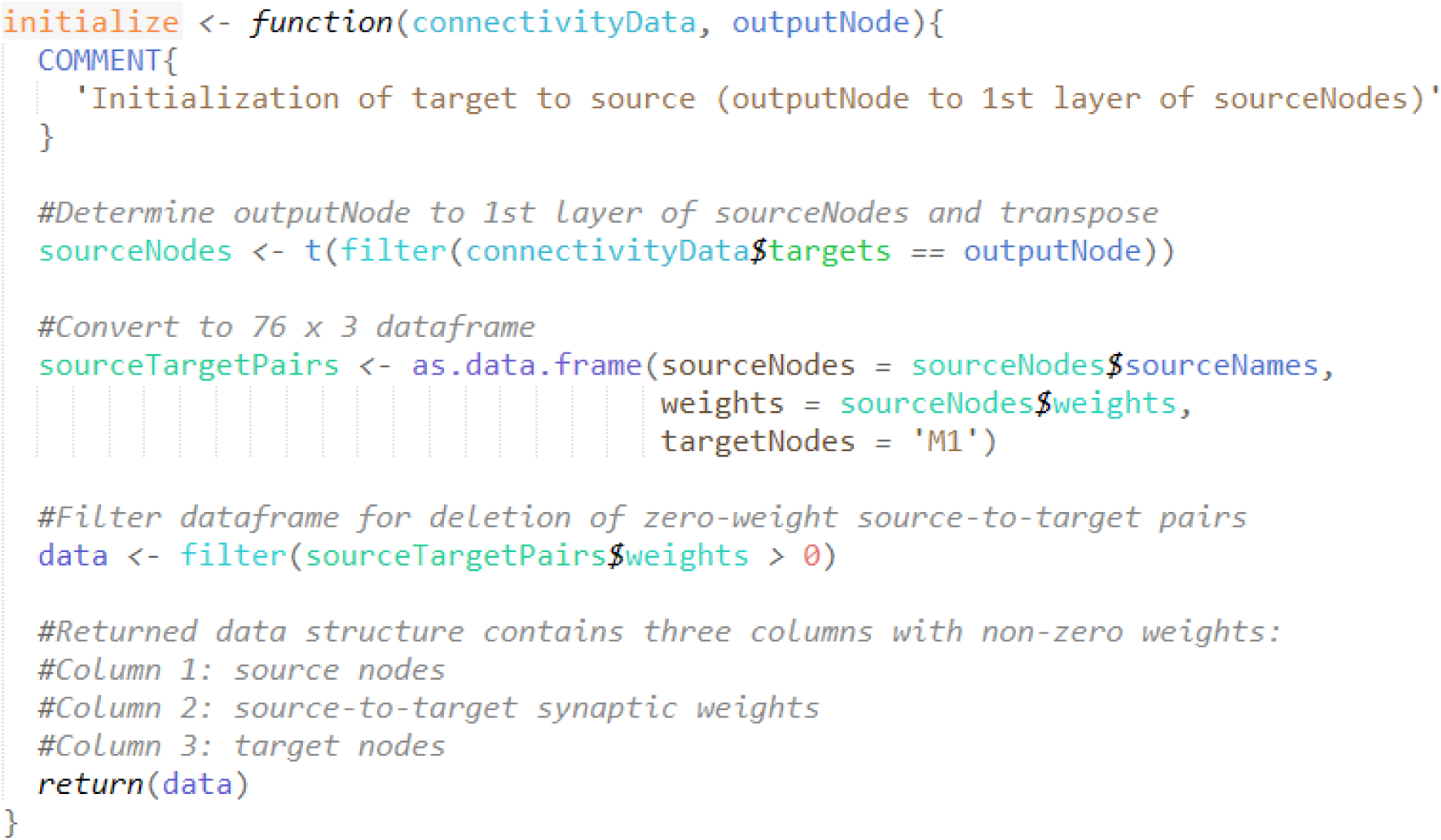
Pseudocode for initializing and finding first layer of regions of the modularity algorithm. This method unpacks the connectome from the output node (M1) and identifies all its source nodes (all brain regions connected to the output node by a single connection, “layer 1”).

**Figure 3.**
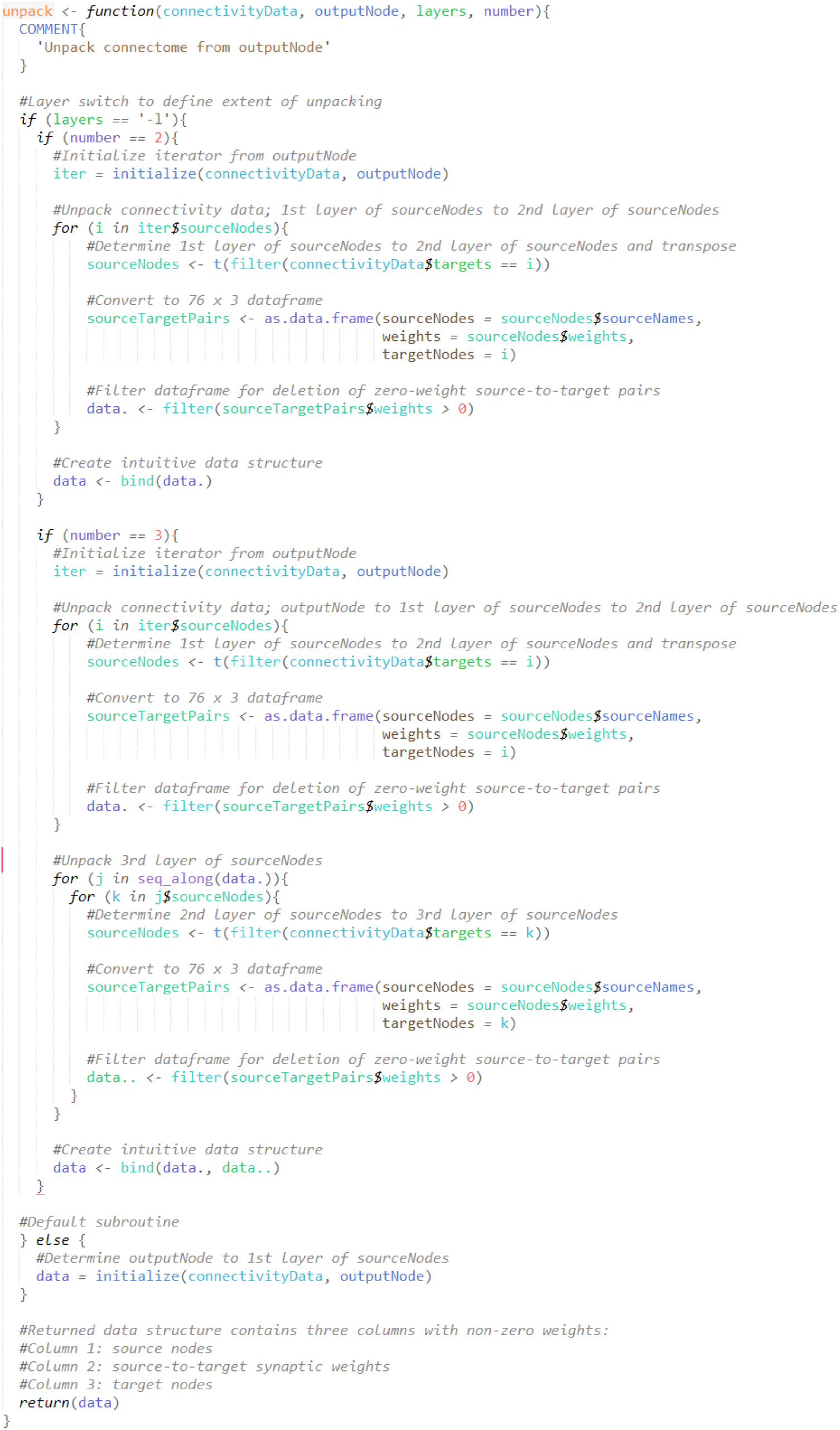
Pseudocode for identifying 2^nd^, 3^rd^, etc. layers from output region, stopping at layer containing input region. This portion unpacks the connectome and finds the regions connected to all the “layer 1” regions identified in Figure 2 by a single connection (“layer 2”), then all the regions connected to these “layer 2” regions by a single connection (“layer 3”), etc. Here, identifying “layer 2” was sufficient to find the input region V1.

**Figure 4.**
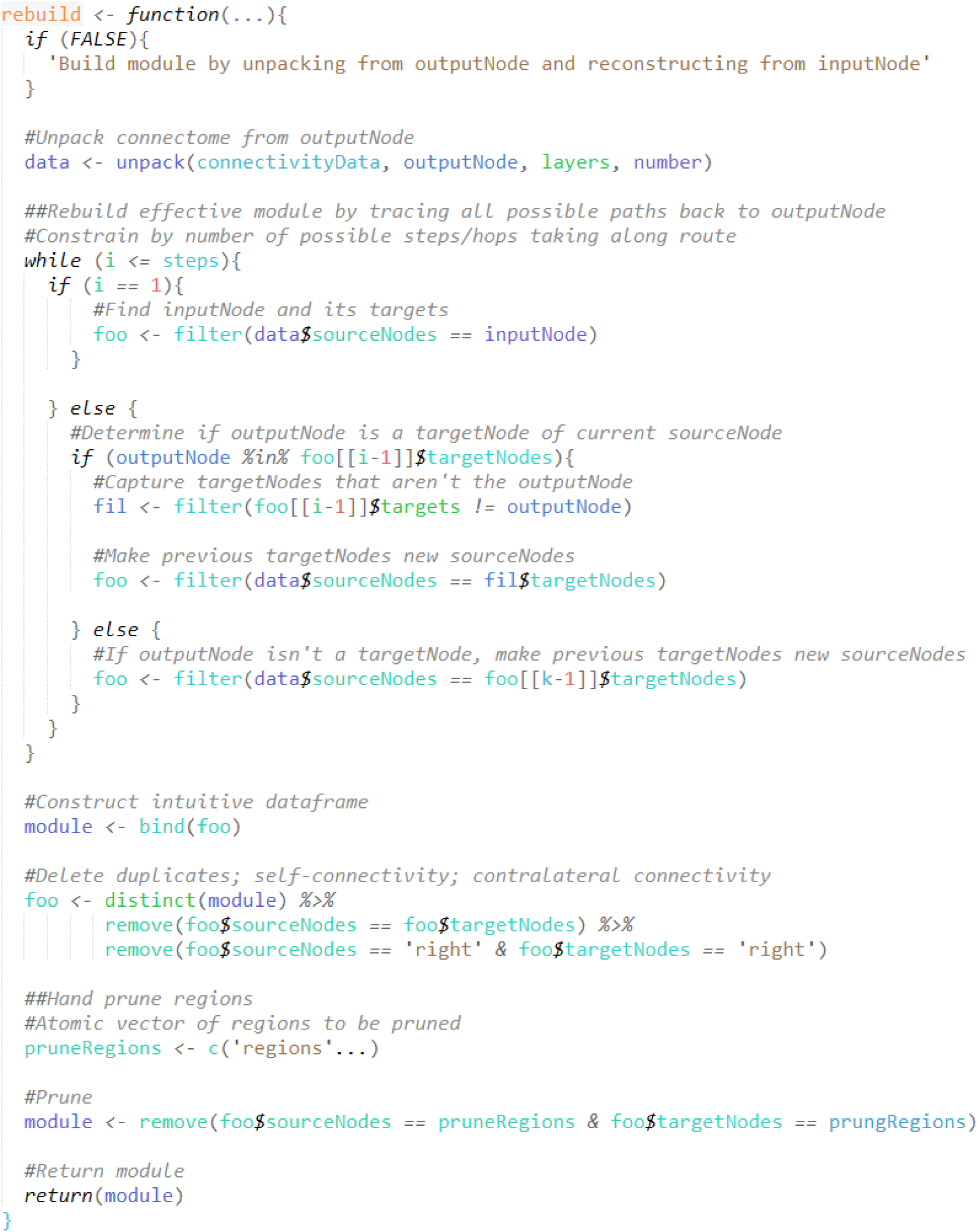
Rebuild pseudocode. This function finds all possible paths from the input (V1) to output (M1) regions in the layers identified in Figure 3. The function is constrained by a user-set number of steps (here, *k* = 20) the algorithm can take along the followed paths.

**Figure 5.**
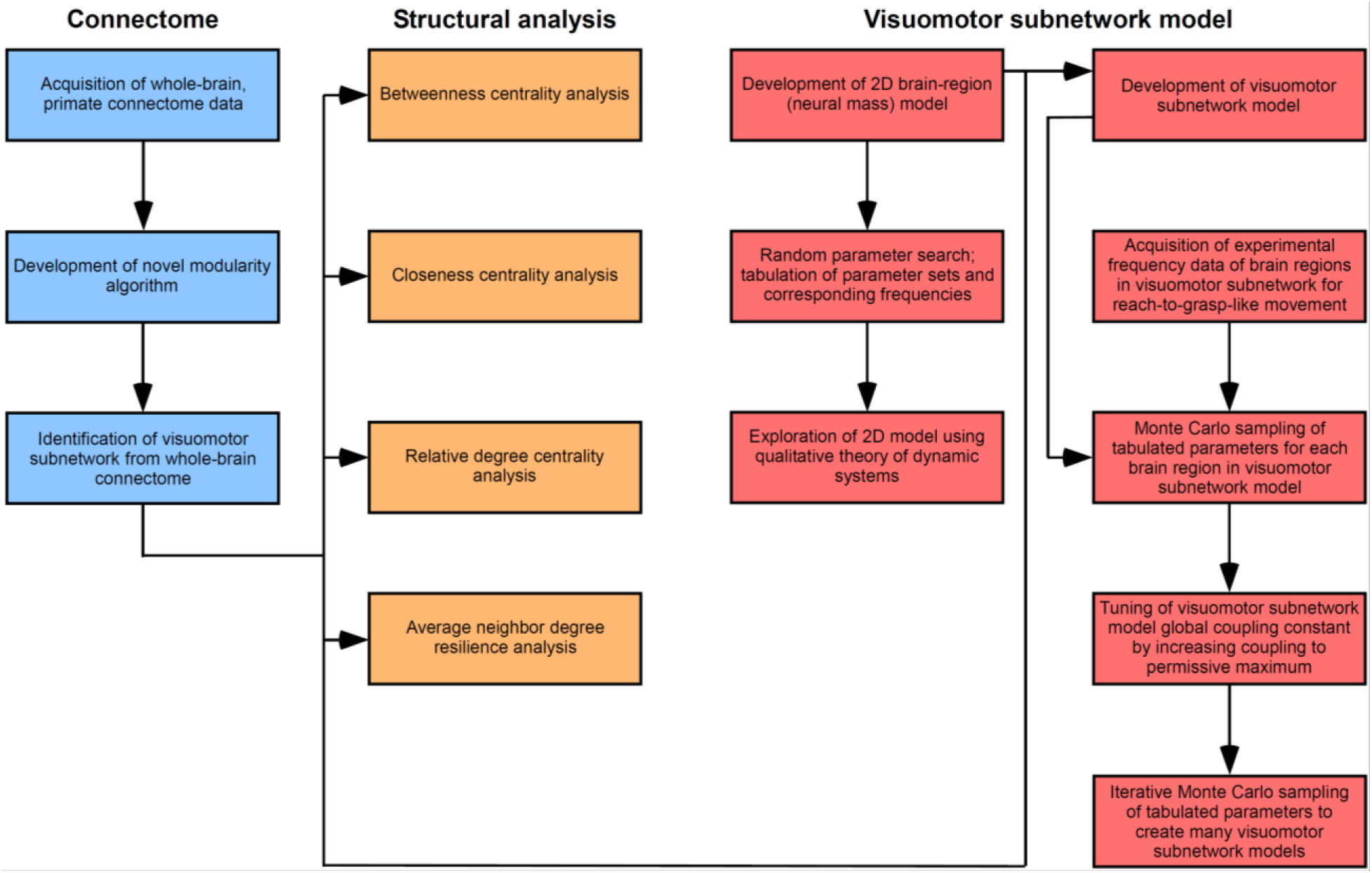
Computational and analytical workflow. The visuomotor functional subnetwork was identified from the brain connectome, graph theory techniques used to identify centrally important regions of the subnetwork and regions particularly likely to alter subnetwork activity, region-appropriate two-dimensional models developed and placed in the subnetwork, and subnetwork and experimental activity compared (columns 1 to 4). Each step involved multiple substeps (entries in each column). Arrows show movement through steps and substeps.

**Figure 6.**
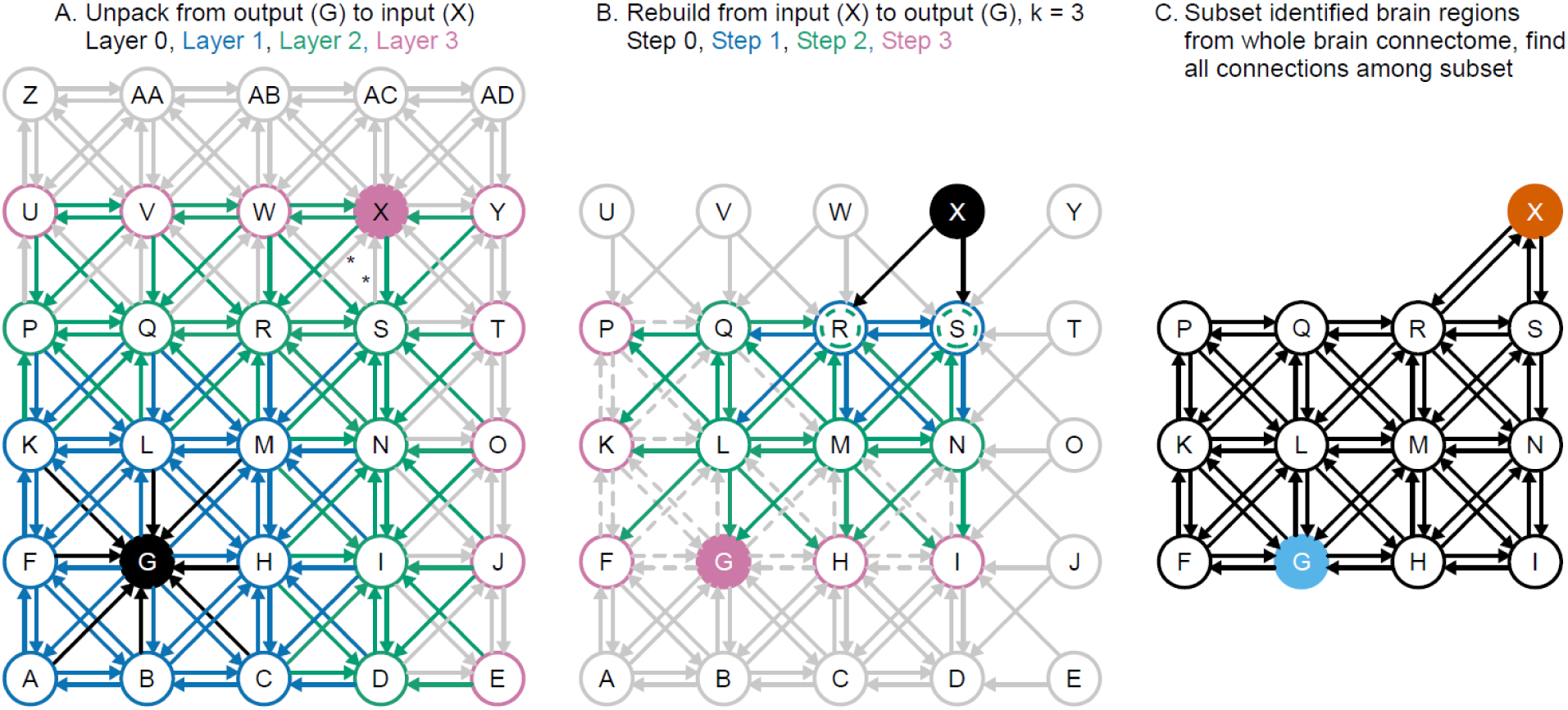
A conceptual framework of the subnetwork identification algorithm. A. Layer identification (unpacking function). The algorithm begins by identifying the region (regions A-C, F, H, and K-L, blue circles) that connect to the output region (G, black arrows). It then finds the regions (blue arrows; regions D, I, N, P-S, green circles) that connect to these regions. It then finds the regions (green arrows; regions E, J, O, T, U-Y, purple circles) that connect to these regions. The process continues until the input region (X) is found. B. Rebuild function. Only regions and connections identified in the unpack function are used. All targets (R and S, blue circles, black arrows) of the input region (X) are found. All targets (Q, L, M, N, green circles, blue arrows) of these regions are then found. Since they reciprocally target each other, the sources R and S are themselves also identified as targets (green dashed circles). The process continues (green arrows, purple circles) for a set number (three here) of iterations. Redundant region identifications due to sources in one step being targets in the next and for other reasons (see text) removed after recursiveness ends. C. The rows and columns of the identified regions are extracted (subsetted) from the tract weight and tract length matrices to construct the final subnetwork.

**Figure 7.**
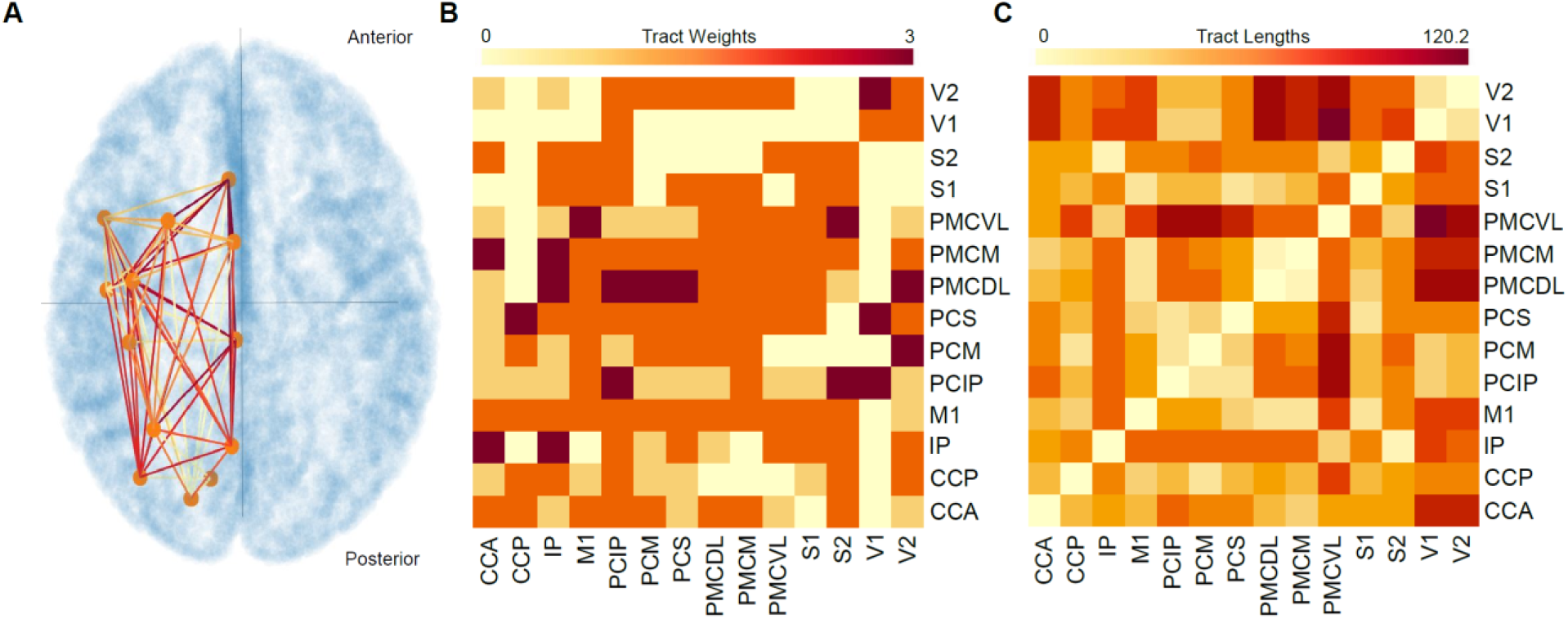
Visuomotor functional subnetwork. **A.** Two-dimensional image of the 3D subnetwork with a representation of cortical surface. Orange nodes are the 14 brain regions present in subnetwork. Orange-to-red edges represent 146 connections between the nodes, different colors used only to help distinguish which edges connect which nodes. Reciprocal and recursive connections not shown for clarity. However, they are included in the module, as can be seen in the tract-synaptic weight matrix in B. Brain orientation: superior-to-inferior along the axial plane, anterior-to-posterior top to bottom. **B.** 14 x 14 square tract-synaptic weight matrix. The matrix is weighted and directed (asymmetric). **C.** 14 x 14 square tract-length matrix. The matrix is weighted but undirected (symmetric).

#### 2.2.1 Finding all regions connected to output region through algorithm initialization (“layer 1” regions; Figure 2)

The first step is using a filter function on the tract-weight matrix to identify all brain regions connected to the output region M1. The required operation is to find all regions that target M1. In the imported data, sources were columns and targets were rows. All rows except the M1 row were therefore deleted. This tensor (in this work, tensor is used as a general term identifying an undefined, heterogeneous data structure like a dataframe) is then transposed, giving a tensor with one column (M1, the target brain region) and 76 rows, the 76 brain regions that potentially connect to M1, with each entry in the column containing the synaptic weight from each region to M1. This tensor is then transformed into a 76 x 3 tensor in which the first column contains the source region of the connection, the second the synaptic weight from this region, and the third the target brain region (in this case, just M1). This 76 x 3 tensor is then filtered for non-zero synaptic-weights, deleting source-to-target pairs with zero synaptic connectivity (i.e., regions that do not connect to M1), which yielded an 18 x 3 tensor.

#### 2.2.2 Finding layers of regions until the layer containing the input region is reached (unpack function; Figure 3)

Subsequent layers of connectivity are found in an analogous manner (Figure 3). Each “layer 1” brain region found in the algorithm initialization is treated in turn as a target region as M1 was treated above when finding the regions that connect to it, resulting in a *n* x 3 tensor for each region. These tensors are then bound row-wise to create a final “layer 1”-“layer 2” tensor with ∑*_r_n_r_* (where *r* refers to which region is being treated as the target) rows and three columns (it was for this reason that the target and source columns were placed in the original tensor). The “layer 2” regions are then used as targets and the process repeated, with a user-set toggle controlling the (integer) number of layers unpacked, until the layer containing the input brain region is found. Here, the input node, V1, was found after unpacking 2 layers of source nodes.

#### 2.2.3 Finding paths from input to output regions (nodes) (rebuild function; Figure 4)

The tensor at the end of the unpacking function contains “extra” brain regions and lacks some connections compared to what will be the final subnetwork. The rebuild function helps address these issues. This function uses the data from the unpacking function as its dataset and traces paths from the input node to the output node. Then, with a control loop constraining the number of possible steps the function can take (max step number = 20 in the work reported here), it filters the tensor to identify the targets of the input node. Note the difference from the unpacking function, which identifies neurons that are the sources of inputs to a given region. The algorithm then uses a Boolean flag to determine if the output region is in this set of regions. If not, the algorithm identifies the target regions of the regions identified as targets in the first step. If the output region is not found in this step, the algorithm uses those targets as the new set of sources, again identifying their targets. This recursive process continues until the output region is found or user-set maximum step number is reached.

#### 2.2.4 Constructing the final subnetwork

At the endpoint of the rebuild function all regions that are part of the functional subnetwork have been identified. These regions (columns and rows) are then extracted (subsetted) from the complete connectome database. This resultant tensors of brain regions, synaptic weights, and tract lengths comprises the subnetwork that is used in all subsequent work.

### 2.3 Graph-theoretic measures for subnetwork analysis

Three measures of network centrality, which identify nodes that, because of their many-node interactions, are well positioned to play a pivotal role in local network activity, were used (Betweenness, Closeness, and Relative Degree). One measure of network resilience, which identifies regions that are particularly well placed to alter gross network activity, Average Neighbor Degree, was used. The R package NetworkToolbox (Christensen, 2018) was used for the centrality measures, and the Python package NetworkX (Barrat et al., 2003; Hagberg et al., 2008) was used for the resilience measure. NetworkToolbox was built specifically to analyze brain network or connectome datasets. NetworkX is a general tool for complex network analysis.

In using these packages care must be taken to ensure that the correct input data structure row and column format is used. In physics, the tensor convention is that sources are columns and targets are rows. In mathematics the standard convention is that sources are rows and targets are columns. NetworkToolbox uses the physics convention and NetworkX the mathematics convention. The equations originally defining the measures (Rubinov and Sporns, 2009) were in the mathematics tensor configuration. The tensors created at the end of Section **2.2.4 Constructing the final subnetwork** were in the physics form. We therefore here present these equations in the physics convention. Because of the different conventions used in NetworkToolBox and NetworkX, the tensors created at the end of Section **2.2.4 Constructing the final subnetwork** were used as input for the NetworkToolbox analyses and the transpose of these tensors were used as input for the NetworkX analysis.

In all equation presented here, notations in the form of *X_ji_* represent the value of the quality being examined (*X*; e.g., synaptic weight) from *j* (the source column) to *i* (the target row).

#### 2.3.1 Betweenness

Betweenness calculates the fraction of all shortest paths that contain a given node in a network for a weighted, directed graph. Thus, nodes interfacing different paths in a network have higher betweenness values. Rubinov and Sporns (2009) use the equation

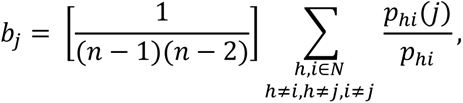

where *b_j_* is the betweenness of node *j*, *p_hi_* the number of shortest paths from *h* to *i*, *p_hi_(j)* the number of shortest paths from *h* to *i* that pass through *j*, *n* the number of nodes, and *N* the set of all nodes.

Network Toolbox uses the same equation without the node-dependent scaling factor, which will not affect the ordering of regions for this measure (see Figure 8A):

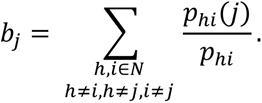

**Figure 8.**
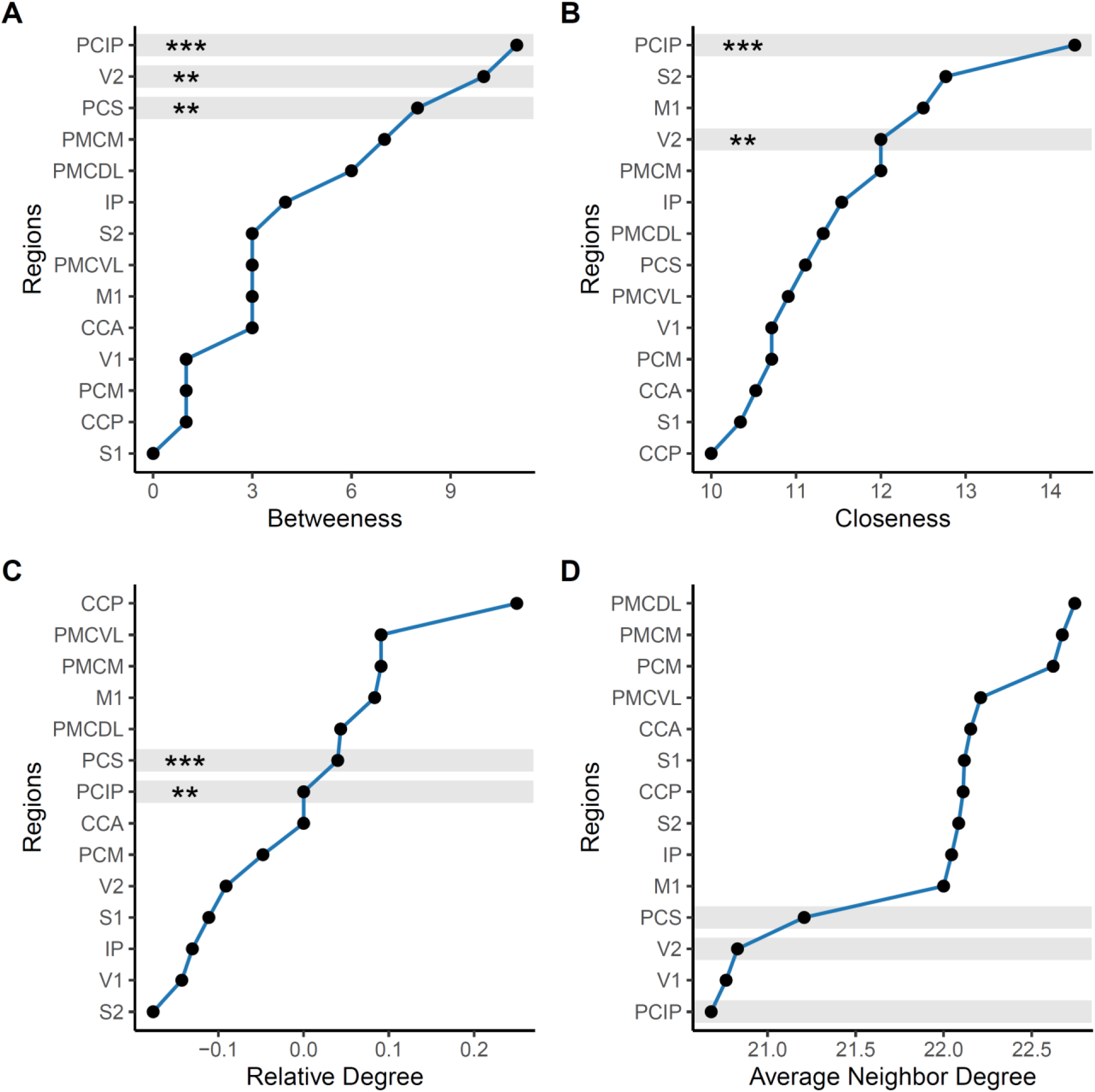
Analysis of visuomotor subnetwork criticalities and resiliency. **A** Betweenness measure of visuomotor subnetwork suggesting PCIP, V2, and PCS are network centralities. **B.** Closeness measure of visuomotor subnetwork suggesting PCIP and V2 are network centralities. **C.** Relative degree measure of visuomotor subnetwork suggesting PCIP and PCS are network centralities. For relative degree, regions with a value at or around 0 were considered the highest scoring regions. In **A**, **B**, and **C**, regions that had centrality measures in the top approximately 25% in at least two metrics are grey highlighted, asterisks indicate in how many centrality measures the region was in the top 25%. **D.** Low average neighbor degree measure of the high centrality regions suggest that PCIP, V2, and PCS are regions in which changes in activity are most likely to change gross visuomotor subnetwork activity.

**Figure 9.**
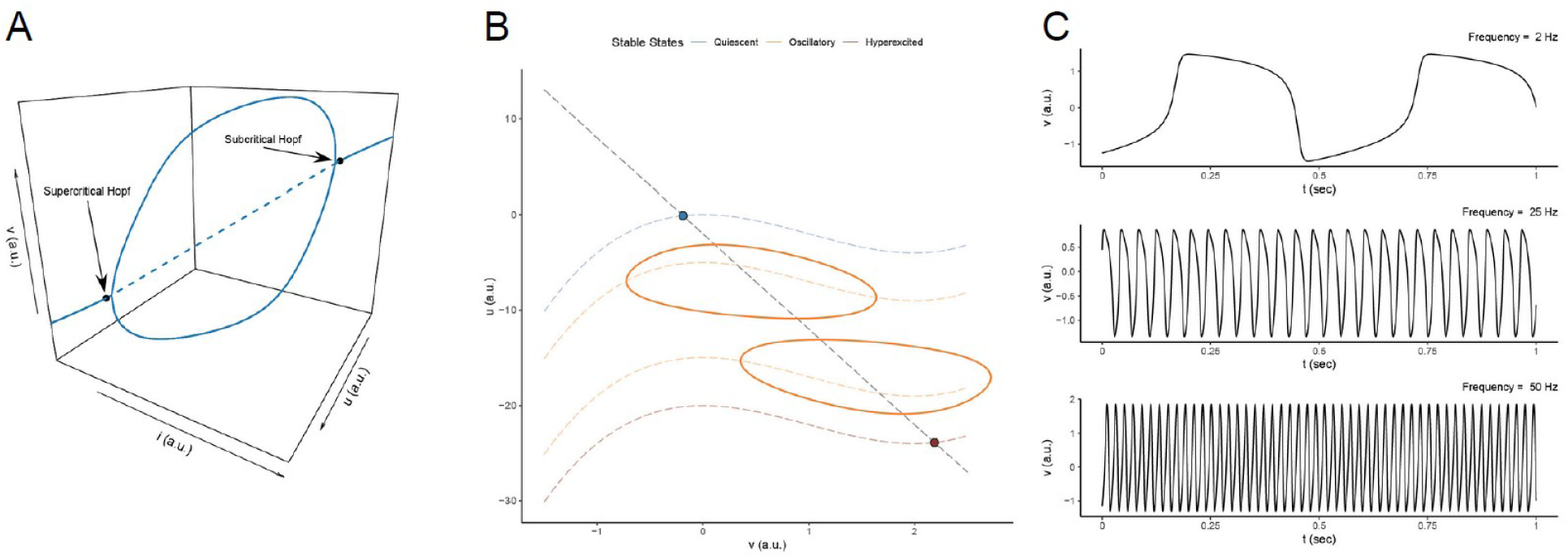
Oscillatory activity, non-linear responses of simplified brain region model (A, B) and models producing different frequency oscillations (C). A. A 3D bifurcation analysis of the steady-state activity of the simplified brain region model. At low *i* the model produced increasingly depolarized single *v* activity. As *i* is further increased the model underwent a supercritical Hopf bifurcation leading to a limit cycle attractor (oscillation as the steady-state activity) and then a subcritical Hopf bifurcation leading again to a stable single *v* attractor. **B**. A 2D phase plane illustrating the primary activity regimes of the brain-region model. The model’s low *v* stable-state activity, oscillatory activity, and high *v* activity result from vertical transpositions of the *v* (cubic) nullcline. **C**. Three of multiple models that oscillate in the 2, 25, and 50 Hz ranges, corresponding to delta, beta, and gamma waves, found by searching across all model parameters.

#### 2.3.2 Closeness

Closeness is a measure of the distance from a given node to all other nodes in the network and is often defined as the inverse of the average shortest path length of a node to every other node (Liu et al., 2017; Rubinov and Sporns, 2009). Thus, the smaller the average path length, the higher the centrality measure because the node is less distant from all other nodes. For a weighted, directed graph, Rubinov and Sporns (2009) use the equation

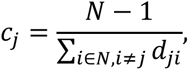

where *c_j_* is the closeness of node *j* and *d_ji_* is the computed shortest path lengths from node *j* to all other nodes *i*. NetworkToolbox uses the same equation without the node-dependent scaling factor, which will not affect the ordering of regions for this measure (see Figure 8B):

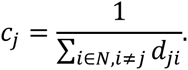

#### 2.3.3 Relative Degree

“Degree” identifies the total number of edges connected to a node irrespective of direction (*Nodes_in_ + Nodes_out_*). “Relative degree” measures the number of outgoing connections relative to the number of incoming connections: (*Nodes_out_ – Nodes_in_*)/(*Nodes_out_ + Nodes_in_*). Positive values indicate more outgoing connections than incoming connections, negative values more incoming than outgoing. For a weighted, directed graph, relative degree is given by the same equation in Rubinov and Sporns (2009) and NetworkToolbox

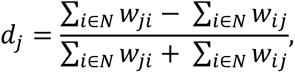

where *w_ji_* is the weight of the interconnection (edge) from *j* to *i* and *w_ij_* the weight of the interconnection (edge) from *i* to *j*.

#### 2.3.4 Average Neighbor Degree

Average neighbor degree is a local assortativity metric that assesses the resilience of overall network function to changes in the activity of its individual components. The lower a brain region’s average neighbor degree, the more likely changes in its activity are to disrupt overall network activity, particularly if the region is also central to the network (Hagberg et al., 2008; Pastor-Satorras and Vespignani, 2001; Rubinov and Sporns, 2009). For a weighted, directed graph, average neighbor degree is given by (NetworkX uses the same)

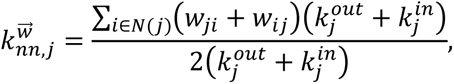

where 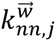 is the average neighbor degree of node *j*, *w_ji_* and *w_ij_* are as defined above, *k_j_^out^* is the out degree for node *j*, and *k_j_^in^* is the in degree for node *j* and the equation refers to the untransposed tensor obtained in **Section 2.2.4 Constructing the final subnetwork**.

For relative degree and average neighbor degree, connection strength is the appropriate connectivity data. These analyses were therefore performed on the visuomotor subnetwork’s 14 x 14 tract-synaptic weight matrix (Figure 2B). For betweenness and closeness, distance is the appropriate connectivity data. However, the algorithms used here handle weight matrices by computing their inverses (Brandes, 2001; Christensen, 2018; Rubinov and Sporns, 2009). Thus, synaptic weights were transformed into lengths by assuming that an increase in weight corresponded to a decrease in distance. As such, the visuomotor subnetwork’s 14 x 14 tract-weights matrix was used in the betweenness and closeness calculations as well.

### 2.4 Implementation details of a generic brain region model (see also exploration of model in Figure 9, Table 3)

All simulations in this section were integrated using a 2^nd^ order Runge-Kutta method and run for times (*t* ≥ 5s) that trial and error showed were sufficient to reach steady-state. All parameters were unitless except time (*t*), which was in ms.

**Table 3.** Parameter sets used in Figure 9.

| parameter | 2 Hz | 25 Hz | 50 Hz | default |
| --- | --- | --- | --- | --- |
| $a$ | -4.54 | 2.51 | -3.52 | -2.00 |
| $b$ | -7.65 | 13.91 | 13.86 | -10.00 |
| $c$ | 4.34 | -0.68 | 0.34 | 0.00 |
| $d$ | 0.0033 | 0.034 | 0.054 | 0.02 |
| $e$ | 0.34 | -3.31 | 3.01 | 3.00 |
| $f$ | 2.52 | 3.24 | 1.26 | 1.00 |
| $g$ | 3.82 | 1.14 | 0.074 | 0.00 |
| $\alpha$ | 3.94 | -2.22 | -2.90 | 1.00 |
| $\beta$ | -3.52 | -3.66 | -1.21 | 1.00 |
| $\gamma$ | 0.24 | 0.44 | 0.25 | 1.00 |
| $i$ | 1.52 | 1.18 | -1.95 | 0.00 |
| $\tau$ | 4.84 | 2.42 | 1.21 | 1.00 |

#### 2.4.1 Simplified model at different constant *i* (see also Figure 9A, 9B)

The simplified model (Table 3 “Default” column except *i* was varied) was simulated with Δ*t* = 0.025 ms. The minimum and maximum of each state variable for each run was determined for each *i*. When *v_min_ = v_max_* and *u_min_ = u_max_*, the model was quiescent (a single {*u*,*v*} was its stable attractor). When *v_min_ ≠ v_max_* and *u_min_ ≠ u_max_*, the steady-state activity was oscillatory. In the top panel of Figure 9B the four *v* nullclines (from top to bottom) are with *i* equal to zero, 5, 15, and 20, respectively.

#### 2.4.2 Finding model parameters that give oscillations in macroscopically relevant frequency bands (same-named columns in Table 3)

Multiple brain region models that produced delta, beta, and gamma frequency oscillations were found by a random parameter search in the ranges of a ∈ (−5, 5), b ∈ (−20, 10), c ∈ (−10, 10), d ∈ (0.0001, 1), e ∈ (−5, 5), f ∈ (−5, 5), g ∈ (−5, 5), α ∈ (−5, 5), β ∈ (−5, 5), γ ∈ (−1, 1), i ∈ (−5, 5), τ ∈ (1, 5), units arbitrary. Each parameter was randomly selected using a floating-point pseudo random number generator. The values for each parameter were uniformly distributed across the range. The brain-region model was then simulated in parallel across n_cores_ – 1 for n_simulations_ = 15,000 runs with a time step Δ*t* = 0.02 ms, chosen to decrease computational cost across n_simulations_. The voltage activity during the last one second of the simulation was then Fourier transformed and the maximum amplitude frequency identified as the dominant frequency. The dominant frequency and corresponding parameter set were saved if the frequency was between 2 and 100 Hz. This search resulted in 13,821 parameter sets.

## 3 RESULTS

The work reported here consisted of four steps: identifying the visuomotor functional subnetwork from the brain connectome, identifying centrally important regions of the subnetwork and regions particularly likely to alter gross subnetwork activity, constructing a 2D model to simulate brain region inherent activity and choosing appropriate versions of the model for each region, and placing these models in the subnetwork and comparing visuomotor model and experimental activity, with each step involving multiple substeps (Figure 5).

### 3.1 Identifying visuomotor functional subnetwork from brain connectome

#### 3.1.1 Neurobiological background

Visual input (light) is transduced at the retina, which communicates via the optic nerve to the lateral geniculate nucleus of the thalamus. Visual information leaves the thalamus and travels to V1, the first cortical vision-related brain region in the primate connectome, via the geniculostriate pathway. For visuomotor behaviors, this input ultimately results in a motor output, and thus ends at the primary motor cortex (M1), where motor initiation occurs (Kandel et al., 2000). The modularity algorithm therefore used V1 as the input region (node) and M1 as the output region (node) in extracting the visuomotor functional subnetwork from the entire brain connectome.

#### 3.1.2 Identifying (unpacking) all layers of regions between output and input region

For generality and ease of visualization, rather than the data shown in Figure 1B, we demonstrate here how the algorithm identifies a functional subnetwork from a connectome using a generic synaptic interconnectivity consisting of six rows of five neurons each, where each region is connected to all its nearest neighbors (Figure 6). Region G was the output region and region X the input region. The first step in the algorithm (Figure 6A; see Figure 2 and associated text in Material and Methods for details) was identifying all regions that made inputs to region G (black arrows pointing toward G). This identified eight “layer 1” regions (A-C, F, H, and K-L; blue circles). Region X was not in this layer. Regions that made inputs (blue arrows) to A-C, F, H, and K-L were therefore identified. Note that it is not until this second step that region G’s output connections are identified (blue arrows from G to neurons A-C, F, H, and K-L). These “layer 2” regions were D, I, N, and P-S (green circles). Region X was not in this layer. The regions that made inputs (green arrows, “layer 3”) to D, I, N, and P-S were therefore identified (E, J, O, T, and U-Y; reddish-purple circles). Region X was in this layer. The unpacking function therefore stopped at this point. Note that, due to inputs to a given layer’s regions not being identified until the step after each layer’s regions had been identified, two connections that are certainly a part of the subnetwork (the grey connections from region R and S to X, asterisks) have not yet been identified.

#### 3.1.3 Identifying (rebuilding) a pathway between the input and output region (node) subject to the connection length restriction

The unpacking function identifies a set of expanding layers centered, as allowed by the topology of the set of brain regions being examined, on the output layer. That is, if in Figure 6A, there were more columns to the left of the leftmost column and more rows below the bottom row, the green and purple regions would form complete rectangles centered about region G. As such, at this stage the only constraint on which brain regions are found that depends on the input region is the number of layers (in this example, three). Although it is certainly possible that functionally important pathways between the output and input regions are present in this large set of brain regions (e.g., G to H to C to D to E to J to O to G to Y to X), this set clearly does not in any sense identify only paths that most centrally connect G and X (e.g., G to M to S to X).

One could decrease the number of seemingly less centrally-connecting pathways by limiting the number of brain regions allowed on any pathway connecting the output and input regions. However, such a procedure would not make the subnetwork selection process more dependent on the input layer and would be completely arbitrary. It seemed to us a better alternative would be a function in which the input region played a role analogous to the role the output region plays in the unpack function. In the unpack function region G serves as the start point, the anchor, for the identification process. Using the input region (X) as the starting point (anchor) in a connection search to the output region (G) through the brain regions identified by the unpack function would thus fulfill the goal of the output and input regions both playing a major role in subnetwork identification.

In the case at hand (Figure 6B), the input to this “rebuild” function is the colored regions and connections in Figure 6A. The first step of the rebuild function is to discard to which layer each region belonged in the unpack function (i.e., all the regions identified by the unpack function become grey). Step two is to identify the targets of the input region (X, black region and arrows) from the set of connections identified by the unpacking function (note that, for example, region W can never be identified in the rebuild function because no input connection to W was identified in the unpack function). These regions are R and S. These regions then become the source regions in step two (blue circles and connections), identifying regions Q and L-N as targets. These are all brain regions that can (from the set of connections available to the rebuild function) be connected to region X by two connections. In this step R and S are also identified as step two targets, since R and S are sources and targets of each other (note that each are also regions that are two connections from X (e.g., X to S and S to R makes R two connections away from X; dashed green small circles in R and S regions). Thus, the entire set of targets identified by the algorithm at the end of step 2 is Q, L-N, and R and S. Regions Q, L-N, R and S now become the step three source regions. The Q and L-N targets identify new brain regions (F-I, K, and P; green circles and connections) that can be connected to region X by three connections.

Just as with the unpack function, this process could continue far beyond the output layer if the connectome topology allowed it (e.g., in the next step identifying here the regions A, B, C, and D and, were the total set of brain regions expanded to include a column to the left of the present leftmost column, regions in this column). Doing so again defeats the goal of finding what seems to be a core or central subnetwork connecting the input and output regions. The method to prevent this “oversearching” is again to stop the search when the target (in the rebuild function, the output region) is found. In this example, this was at the end of the third step, but the program allows the number of rebuild steps to be set by the user. In all work reported here the rebuild steps were set to the number necessary to find the output (M1) layer.

With respect to the data stored, in addition to the single instances of the F-I, K, and P being identified in step 3, there are also redundancies due to, in the specific example given above, R and S being sources at the beginning of both Steps 2 and 3. This leads to R and S being identified in step 1 because they are targets of X, re-identified in step 2 because they are targets of each other, and then again re-identified in step 3 because they are targets of each other and of regions L-N and Q, all of which instances are present in the regions that have been stored to now in the rebuild function.

When applied to the real brain connectome, right brain regions due to callosal connections were also present, which should not be because of left-brain visuomotor lateralization (Frey et al., 2005; Gonzalez et al., 2006; Meador et al., 1999; Perenin and Vighetto, 1988; Radoeva et al., 2005; Smutok et al., 1989). The visual system also contains two components, a posterotemporal ventral stream responsible for “what” driven, perceptual object identification and a posterofrontal dorsal stream responsible for analyzing “where” based spatial information necessary for action (Goodale and Milner, 1992; Sheth and Young, 2016; Ungerleider and Mishkin, 1982), albeit with substantial crosstalk between the two components (Budisavljevic et al., 2018; van Polanen and Davare, 2015). For reach-to-grasp-like movements we were only interested in the “where”-involved regions. Cross-referencing the brain regions initially identified in the rebuild function with respect to being involved in motor, visual, and visuomotor function (Friedrich, 1933; Nelissen and Vanduffel, 2011), as well as descriptions of their functions from the literature (Berens et al., 2008; Frien et al., 1994; MacKay and Mendonca, 1995; Murthy and Fetz, 1992; Rougeul et al., 1979; Sanes and Donoghue, 1993; Stetson and Andersen, 2014; Weiss et al., 2018; Womelsdorf et al., 2010), identified four regions to be removed from the subnetwork on the basis of not likely being involved in reach-to-grasp movement production (Table 2).

Redundant, callosal, and inappropriate regions were all removed from the network identified at the end of the rebuild function (end of Figure 4 pseudocode).

#### 3.1.4 Identifying connections among subnetwork neurons

At the end of the rebuild function, a large percentage of the connections among the subnetwork regions have not been identified (Figure 6B, grey dashed connections). The rebuild function therefore outputs only the identities of the subnetwork regions. These data are then used to create a subset of the entire brain connectivity matrices (tract weight, tract length) by deleting all rows and columns not identified by the rebuild function from the entire brain connectome dataset. This set of regions and their interconnections (all of which are necessarily included, since they are present in the entire brain connectome matrices) are the identified functional subnetwork.

#### 3.1.5 Final visuomotor functional subnetwork (Table 2, Figure 7)

The final visuomotor functional subnetwork contained 14 brain regions (Table 2) and 146 interconnections, all in the left hemisphere (Figure 7A). This restriction to a single hemisphere changed the properties of the tract-synaptic weight (Figure 7B) and tract-length (Figure 7C) matrices. The tract-synaptic weight matrix became a weighted, directed (asymmetric; V1 to V2 synaptic weight is not necessarily the same as V2 to V1 synaptic weight) graph. The tract-length matrix, alternatively, became a weighted, undirected (symmetrical) graph because the distance between any two regions in a single hemisphere is a single number (e.g., the distance between V1 to V2 and V2 to V1 is the same).

### 3.2 Centrally important regions of the subnetwork, regions particularly likely to alter subnetwork activity

#### 3.2.1 Centrally important regions of the subnetwork

Centrally important brain regions were identified on the basis of being in the top four regions (approximately 25% of 14 brain regions in the functional visuomotor subnetwork) in at least two of the three (betweenness, Figure 8A; closeness, Figure 8B; relative degree, Figure 8C) centrality measures. For betweenness and closeness this means regions with the highest absolute values. For relative degree, the highest scoring regions were those at or near 0, as these nodes have equal numbers of incoming and outgoing connections, and, therefore, have equal chance to be perturbed by incoming connections or perturb downstream regions through outgoing connections.

PCIP had the highest measure across each centrality metric (grey in Figures 8A, 8B, and 8C; asterisks indicate the number of occurrences across all centrality measures). PCIP is thus the “primary hub” of the visuomotor subnetwork as it participates in the most intra-subnetwork paths, is in closest proximity to all other regions in the subnetwork, and interacts with the most regions across the visuomotor subnetwork. Its centrality is further supported by the suggestion that PCIP interfaces the visual and motor systems (Nakamura et al., 2001) and functions as an integration area for visual-to-motor transformation (Grefkes and Fink, 2005).

V2 scored in the top 25% in betweenness and closeness (grey in Figures 8A and 8B), making it a “secondary hub” of the functional visuomotor subnetwork. This identification is consistent with V2 being often considered the last pure visual region before the visual pathway diverges into its ventral and dorsal streams (Sheth and Young, 2016) dedicated, respectively, to object identification and vision for action. PCS also scored in the top 25% in betweenness and relative degree (grey in Figures 8A and Figure 8C), identifying it as another “secondary hub” of the functional visuomotor subnetwork, having a presence in many of the intra-subnetwork pathways and relatively high-density connections with other regions in the visuomotor subnetwork.

#### 3.2.2 Regions particularly likely to alter subnetwork activity

The primary and secondary centrality hubs were then compared to the 25% of nodes scoring lowest on the average neighbor degree resilience measure, as these nodes have the highest potential of changing gross subnetwork activity. PCIP, the primary hub of the visuomotor subnetwork, had the lowest average neighbor degree. The secondary hubs, V2 and PCS, had the third and fourth lowest resilience metric, respectively. Thus, changes in PCIP, V2, and PCS activity are most likely to alter visuomotor functional subnetwork activity.

### 3.3 A two-dimensional model that produces a wide range of outputs

#### 3.3.1 The generic brain region model

As noted in **Section 1 Introduction**, when active as a part of performing a task, neurons in a brain region tend to oscillate in phase and are thus often modeled as a unit (Jansen and Rit, 1995; Leon et al., 2015; Stefanescu and Jirsa, 2008; 2011; Wilson and Cowan, 1972; 1973; Wong and Wang, 2006). We modeled here the activity of each brain region as a unit using a widely-used 2D model (Leon et al. (2015):

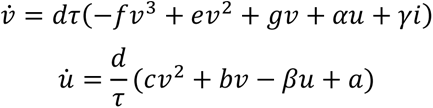

where *v* is a state variable representing the average population voltage activity, *u* represents a relatively slowly resetting state variable, *i* is a variable representing region excitation, and *d*, τ, *f*, *e*, *g*, α, γ, *c*, *b*, *β* and *a* are parameters that can freely vary to produce different model outputs. A detailed description of each parameter can be found in (Leon et al., 2015).

#### 3.3.2 The model can produce multiple steady-state oscillatory outputs

The model can produce a very wide range of outputs as *i* or its free parameters are varied. In the work reported here, we examine model activity only at steady state. To aid in visualization and ease of understanding, in explaining model activity we used a simplified version in which *f*, *τ*, *α*, *β*, and *γ* were 1; *e* was 3, *a* was -2, *b* was -10, *d* was 0.02; and *c* and *g* were zero (thus turning *u̇* into a linear equation; Table 3, “Default” column) and changed only *i* from 0 to 20 at Δ*i* = 0.1 (Figure 9).

Figure 9A shows these data in a 3D bifurcation plot. In this plot, if model output is a constant voltage, this voltage is plotted. If the model’s output is oscillatory, the minimum and maximum voltages of the oscillation are plotted. The transitions from stable to oscillatory activity and back (arrows) are the bifurcations. At all *i* values, if the initial *v* and *u* values of the model differed from those present in Figure 9A (i.e., the simulation began at a point not on the lines in the figure), model activity would move to the appropriate line(s) (either to the single voltage output or the oscillatory voltage output appropriate for that *i*).

At low *i* values, the steady-state output of the model was a single value that increased with *i* (Figure 9A; initial single line). As *i* increased, the model reached a criticality (downward arrow) and underwent a supercritical Hopf bifurcation (Izhikevich, 2007) and became oscillatory (two lines in Figure 9A; in dynamic systems jargon, a limit cycle attractor appeared). Oscillation amplitude (the distance between the two lines in Figure 9A) initially increased with increasing *i*. As *i* continued to increase, oscillation amplitude began to decrease and eventually (rightward arrow) model output again became a single voltage (another bifurcation, in this case a subcritical Hopf bifurcation (Izhikevich, 2007)).

These activities can be understood by examining the system’s nullclines, the *u* and *v* values at which *v̇* (the *v̇* nullcline) or *u̇* (the *u̇* nullcline) equal zero, in a phase space plot (Figure 9B; (Izhikevich, 2007; Mira, 1997)). With the parameter values used here, the *u̇* nullcline is linear (black straight dashed line in Figure 9B). *u̇* is positive below/to the left of the *u̇* nullcline. When the system is in this portion of the phase plane, *u* therefore increases with time. When the system is in the portion of the phase plane above/to the right of the *u̇* nullcline, *u̇* is negative and *u* therefore decreases with time. *u̇* does not depend on *i*, and the *u̇* nullcline therefore does not change when *i* is changed.

With the parameter values used here, the *v̇* nullcline is cubic (curved dashed lines in Fig. 9B). *v̇* is positive above the *v* nullcline. When the system is in this portion of the phase plane, *v* therefore increases with time. When the system is in the portion of the phase plane below the *v̇* nullcline, *v̇* is negative and *v* therefore decreases with time. Unlike the *u̇* nullcline, the *v̇* nullcline depends on *i*, shifting downward as *i* increases (the uppermost *v̇* nullcline is with *i* equal zero and the lower ones with increasing *i*).

Systems of this general form have been extremely well studied. The general rule is that, if the intersection of the *v̇* and *u̇* nullclines occurs on an arm of the *v̇* nullcline with positive slope, the intersection is a positive attractor, and model activity will drive (albeit in many cases as a damped oscillator) to the intersection from any initial *u* and *v*. This is the case for the top and bottom *v̇* nullclines, which are on the single-lined portions of the bifurcation diagram in panel A. Alternatively, when the intersection of the *v̇* and *u̇* nullclines occurs on an arm of the *v̇* nullcline with negative slope, the intersection becomes an unstable point, because, if the initial *v* and *u* are not on the intersection, the system does not drive to the intersection. This shift can be understood by noting that when the intersection is on the arms of the *v̇* nullcline with a positive slope, *v̇* “points” back to the intersection if the system is moved horizontally from the intersection, whereas when the intersection is on the arm of the *v̇* nullcline with a negative slope, *v̇* “points” away from the intersection if the system is moved horizontally from the intersection. The stable attractor in this case is instead a limit cycle, resulting in oscillatory activity (orange elliptical shapes in panel B, double-lined portion in panel A). As the intersection of the two nullclines shifts on the negative arm when the *v* nullcline moves up and down, the amplitude and shape of the stable oscillation changes.

#### 3.3.3 Identification of brain region models with delta (2 Hz), beta (25 Hz), and gamma (50 Hz) oscillation frequencies

The ability of this simple, 2D model to produce different period oscillations is one of the reasons it is so widely used, as it allows a single model to reproduce the wide range of oscillatory frequencies observed in LFP recordings of brain regions. The brain regions identified as part of the visuomotor functional subnetwork (Table 2, Figure 7) oscillate at multiple frequencies. It was therefore necessary to find brain region models that did so, and to find multiple at each frequency to test whether subnetwork output depended on brain region model. We therefore generated many brain region models by randomly altering all parameters in the *v̇* and *u̇* equations and identifying models active at the appropriate frequencies (see Materials and Methods). Examples of the generic brain region model’s ability to produce activity in different frequency bands are shown in Figure 9C and the corresponding parameter sets in Table 3.

### 3.4 Incorporating brain region models into the visuomotor functional subnetwork

#### 3.4.1 Linking brain region activities

Having a brain region model allows the activity of one region to alter the activity of another through an additive connectivity equation of the region’s *v̇* equation. This was accomplished with the linear coupling term (Leon et al., 2015):

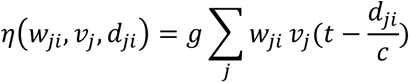

where *g* is the global coupling parameter, w*_ji_* the weight of the connection from the *j^th^* source region to the *i^th^* target region from the synaptic weight matrix of the functional subnetwork, and *v_j_* the voltage of the *j^th^* source region time delayed according to the distance (obtained from the functional subnetwork’s distance matrix) between the two regions (i.e., 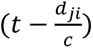 is not a multiplicative term, but indicates that *v_j_* is a function not of the present time, but time delayed by 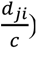. *c* is conduction velocity and was set at 5 m/s, a physiologically realistic value for an adult primate (Firmin et al., 2014; Ghosh et al., 2008; Ivanov and Calabrese, 2006). Each brain region’s *v̇* therefore became the original *v̇* plus *η*(*w_ji_,v_j_,d_ji_*) (the neural mass equation):

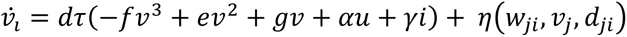

#### 3.4.2 Choosing brain region inherent activities

For 11 of the 14 brain regions in the visuomotor functional subnetwork, dominant oscillatory frequency or frequency range data during monkey and human reach-to-grasp-like movements have been obtained from macroscopic electrophysiological data (Berens et al., 2008; Frien et al., 1994; MacKay and Mendonca, 1995; Murthy and Fetz, 1992; Rougeul et al., 1979; Sanes and Donoghue, 1993; Stetson and Andersen, 2014; Weiss et al., 2018; Womelsdorf et al., 2010). For the three remaining regions (posterior cingulate cortex, posterior insular cortex, and secondary somatosensory cortex), frequency data of macroscopic electrophysiological measurements were used (Olson et al., 1993; Ray et al., 2008; Smuda et al., 2019). These data (each brain region’s oscillatory frequency, *f*) were then used as targets to choose from the parameter-set database built from a large random parameter search described in Sections **2.4.2 Finding model parameters that give oscillations in macroscopically relevant frequency bands** and **3.3.3 Identification of brain region models with delta (2 Hz), beta (25 Hz), and gamma (50 Hz) oscillation frequencies** above to find brain region models that oscillated with a period ± 3 Hz of each region’s dominant frequency during reach-to-grasp-like movements. Multiple brain region models were always found that met this criterion. 30 visuomotor models were then randomly constructed by performing Monte Carlo sampling from the parameter-set database for each brain region model within it (i.e., 14 generic brain region models) that met the brain region in question’s *f* ± 3 Hz criterion. *f*’s of 21 Hz frequency were not allowed as this *f* often produced instabilities in visuomotor model activity, likely due to integration error of the stiff system.

Brain regions are often strongly coupled (Choi and Mihalas, 2019; Cocchi et al., 2017), especially during conscious, focused cognition, or behavior. The global coupling parameter *g* of the connectivity equation was set to the highest value, 0.0055, at which the network continued to produce oscillatory activity. The models were simulated for *t* = 5 seconds at a time step Δ*t* = 0.00005 ms with a 4^th^-order Runge-Kutta integrator to ensure integration accuracy and stability.

### 3.5 Comparison of functional visuomotor subnetwork and experimental activity

We first analyze in some detail one of the 30 visuomotor models selected by the Monto Carlo sampling (Figures 10, 11). Experimental data show that eight brain regions in the visuomotor functional subnetwork produce LFP or LFP-related brain activity in the beta range (12 to 35 Hz), four in the gamma range (>35 Hz), and two in the delta, theta, and alpha ranges (0.5 to 12 Hz). Figure 10A shows the experimental data (black dashed lines, circles), the inherent oscillatory frequency of each of the brain regions (dark red asterisks), and the activity of each region when the subnetwork was interconnected (blue X’s). As expected from coupled oscillator theory, connecting the brain regions altered the oscillation frequencies of most brain regions, bringing some (PCS, PCM, M1) to cycle with the same frequency as another region, PCIP. However, in this instantiation these changes were never large enough to move a brain region’s activity from its inherent frequency band.

**Figure 10.**
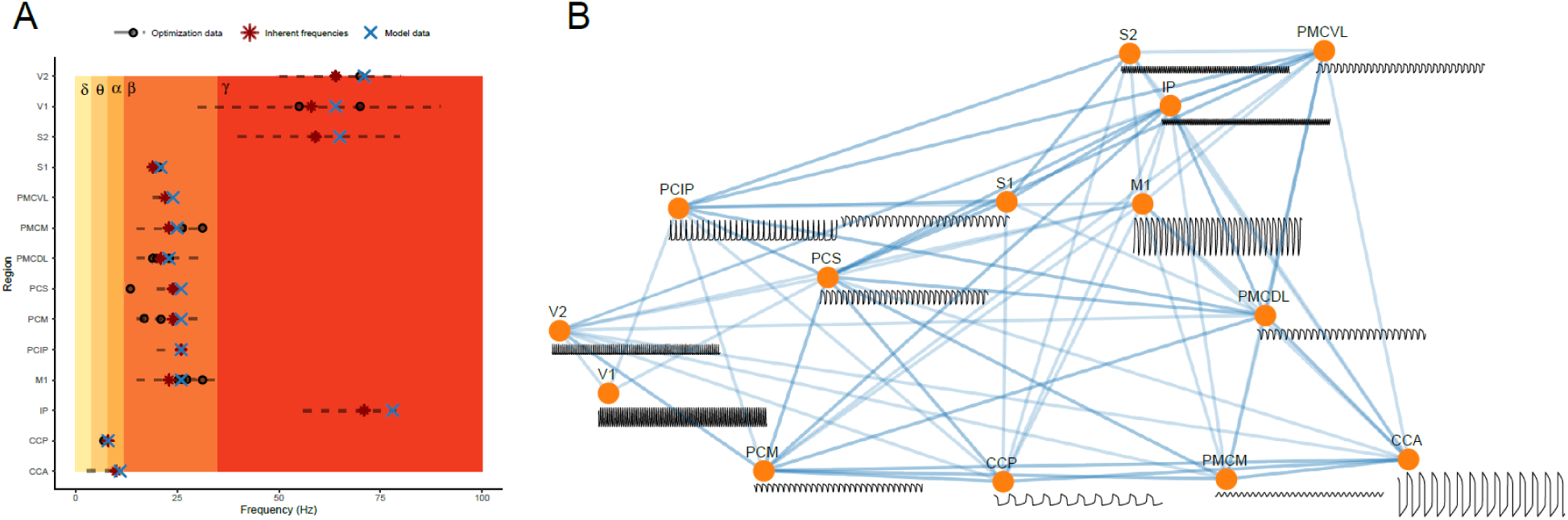
Values and activity of one of the 30 Monte Carlo selected functional visuomotor modules. **A.** Black dotted lines, dots: frequency ranges and dominant frequencies, respectively, reported in the literature. Dark red asterisks: inherent (unconnected to functional visuomotor module) frequency of each brain region. Blue X’s: oscillation frequency of each region in functional visuomotor subnetwork. Interconnecting the brain regions changed the oscillation frequency of all but one region, but all continued to oscillate in their inherent frequency band. Simulation performed with global coupling constant *g = 0.*0055. **B.** A 2D representation of functional visuomotor subnetwork and time-series output of each brain region when in the interconnected subnetwork.

Figure 10B shows a 2D representation of the visuomotor functional subnetwork and the time series activity of each brain region. Amplitude differences in the activity are not meaningful as, experimentally, they depend on the distance of the recording electrode from the brain region and, computationally, arise as a consequence of the frequency-parameter set generation process. The frequency differences, and match between experimental and visuomotor model frequencies in each visuomotor model, alternatively, do demonstrate that the functional visuomotor subnetwork, with this set of brain region models, produces frequency outputs similar to those observed experimentally.

Weakly-coupled oscillator theory predicts that, when oscillators of different inherent frequencies are coupled, the activity of the oscillators should change. In a network with an interconnectivity pattern as complicated as that in Figure 10B, and a range of inherent frequencies as great as that present in the subnetwork regions, a wide range of coupling types (e.g., 1:1, n:m) is expected. This issue is made more complicated by most brain regions being connected to regions with different oscillation frequencies. For example, CCA, with an inherent cycle frequency of about 10 Hz, is connected to CCP, with an inherent cycle frequency of about 8 Hz; PMCM, PCM, PCS, PMCDL, M1, and PMCVL, each with inherent cycle frequencies of about 25 Hz; and V2, IP, and S2, with inherent frequencies between 65 and 80 Hz. Moreover, the effective connection strength varies widely across regions, both with respect to the values in the tract-weight matrix and region oscillation amplitude (which the tract-weight matrix only scales). No easy predictions can therefore be made from weakly coupled oscillator theory about what types of coupling would be expected among the multiple oscillators.

We therefore instead produced Lissajous (1857) (also called Bowditch (Bowditch, 1815)) curves by plotting, time point by time point, the voltage of one region against the voltage of another (Figure 11). Such plots produce curves of varying complexity depending on whether two oscillators are coupled at all and, if coupled, on the type of coupling (1:1, X:1 with X an integer, n:m; here, for simplicity, we refer to both X:1 and n:m cases as n:m). We show here examples of 1:1 coupling (Figure 11A), 2:1 coupling (Figure 11B), and no direct coupling between two regions with very close cycle frequencies (Figure 11C).

**Figure 11.**
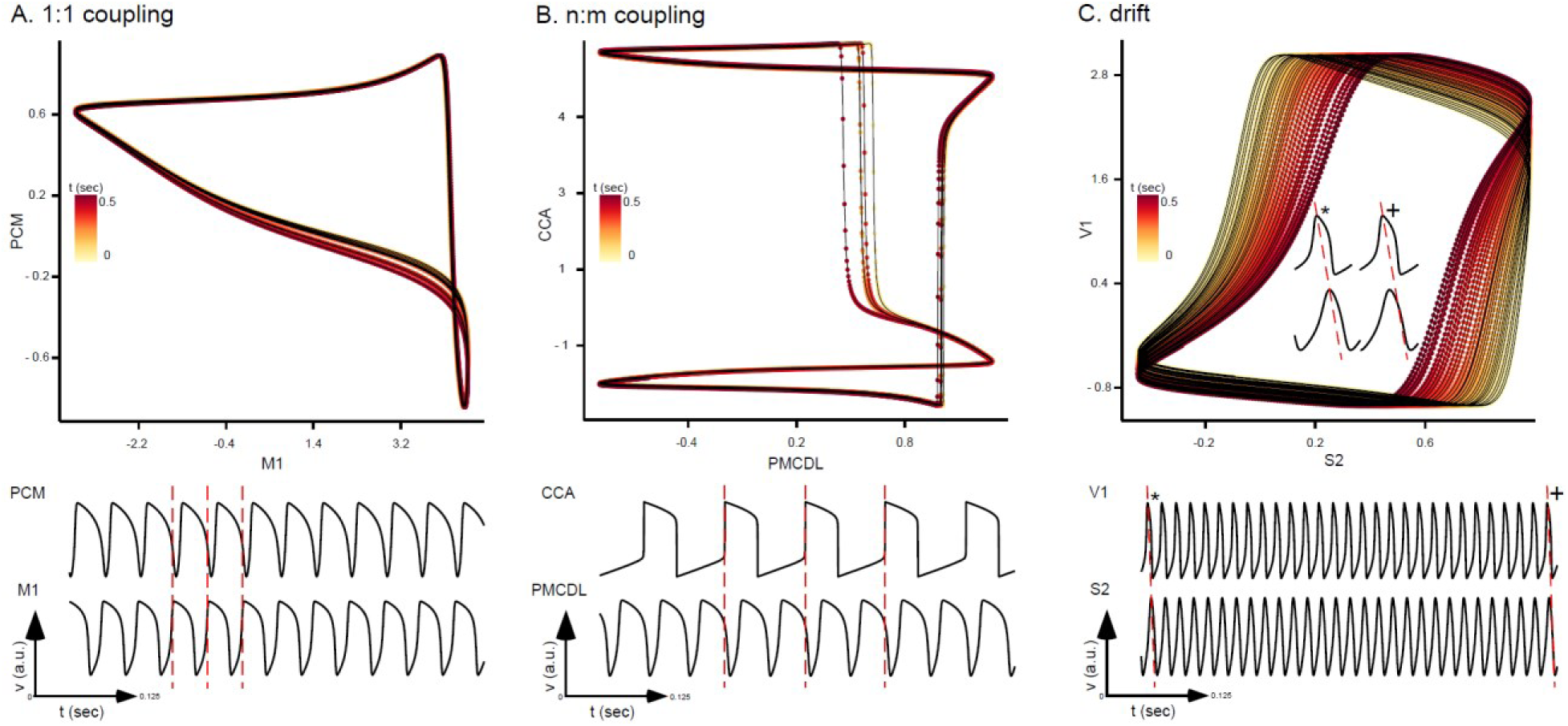
When in the network, the activities of some brain regions entrained (A, 1:1; B, 2:1). The activities of others, even ones with very similar oscillation frequencies, instead continuously drifted relative to one another (C). Upper portion of each panel, Lissajous plot of one brain region’s *v* value vs. another region’s *v* value. Entrained oscillators (A, B) have a single shape that is continuously re-traced as time progresses (yellow to red). Drift is shown by continuously shifting cycles over time (C). Bottom portions of each panel, activity of the two regions shown in each panel over the durations used to produce the Lissajous plots. Entrainment regime (1:1, 2:1) indicated by vertical red dashed lines in A and B. Vertical red dashed lines in C indicate how, by end of 28 oscillations shown, the activity of the two regions has shifted (expansion of 1^st^ (*) and 28^th^ (+) oscillations shown in inset in upper portion of panel; in each case slope of dashed red line identical.)

PCM and M1 have inherent, but different, cycle frequencies of about 25 Hz and are directly connected in the subnetwork. When in the network, the cycle period of both regions slightly increased and the two became 1:1 coupled (Figure 1A). CCA has an inherent frequency of about 10 Hz and PMCDL of about 25 Hz. The two regions are again directly connected in the network. When in the network, the cycle frequency of both brain regions slightly increased from their inherent frequencies (see Figure 10A), with PMCDL’s cycle frequency becoming exactly half that of CCA, and the activity of the two regions entraining 2:1 (Figure 1B). V1 and S2 have inherent frequencies of around 65 Hz. The cycle frequencies of both regions increased from their inherent frequencies when in the subnetwork but did not increase to the same frequency. The two regions are also not directly connected to each other in the network, the most direct connection being through PCIP, which has a very different cycle period (about 25 Hz). The closeness of the cycle periods makes it seem, on first examination, that the two regions are 1:1 coupled. However, the Lissajous plot showed the activity of the two regions actually slowly and continuously drifted through each other, with the peak of S2 activity occurring noticeably earlier relative to the peak of V1 activity at the end of the 28 cycles shown in Figure 11C compared to their relationship at the beginning (inset Figure 11C).

#### 3.5.1 Across-instantiation comparison

As described in Sections **2.4.2 Finding model parameters that give oscillations in macroscopically relevant frequency bands** and **3.3.3 Identification of brain region models with delta (2 Hz), beta (25 Hz), and gamma (50 Hz) oscillation frequencies**, many brain region models (parameters in the *v̇* and *u̇* equations) were found that produced oscillation frequencies near each brain region’s experimental oscillation frequency. The Monte-Carlo sampling chose randomly, from the parameter-set database, parameters such that the *v̇*, *u̇* equations for any given region were different, resulting in 30 different visuomotor models. It was therefore important to examine whether the ability of the visuomotor model to reproduce the experimental data shown in Figure 10 depended on the *v̇*, *u̇* pairs present in each model instantiation (Figure 12).

**Figure 12.**
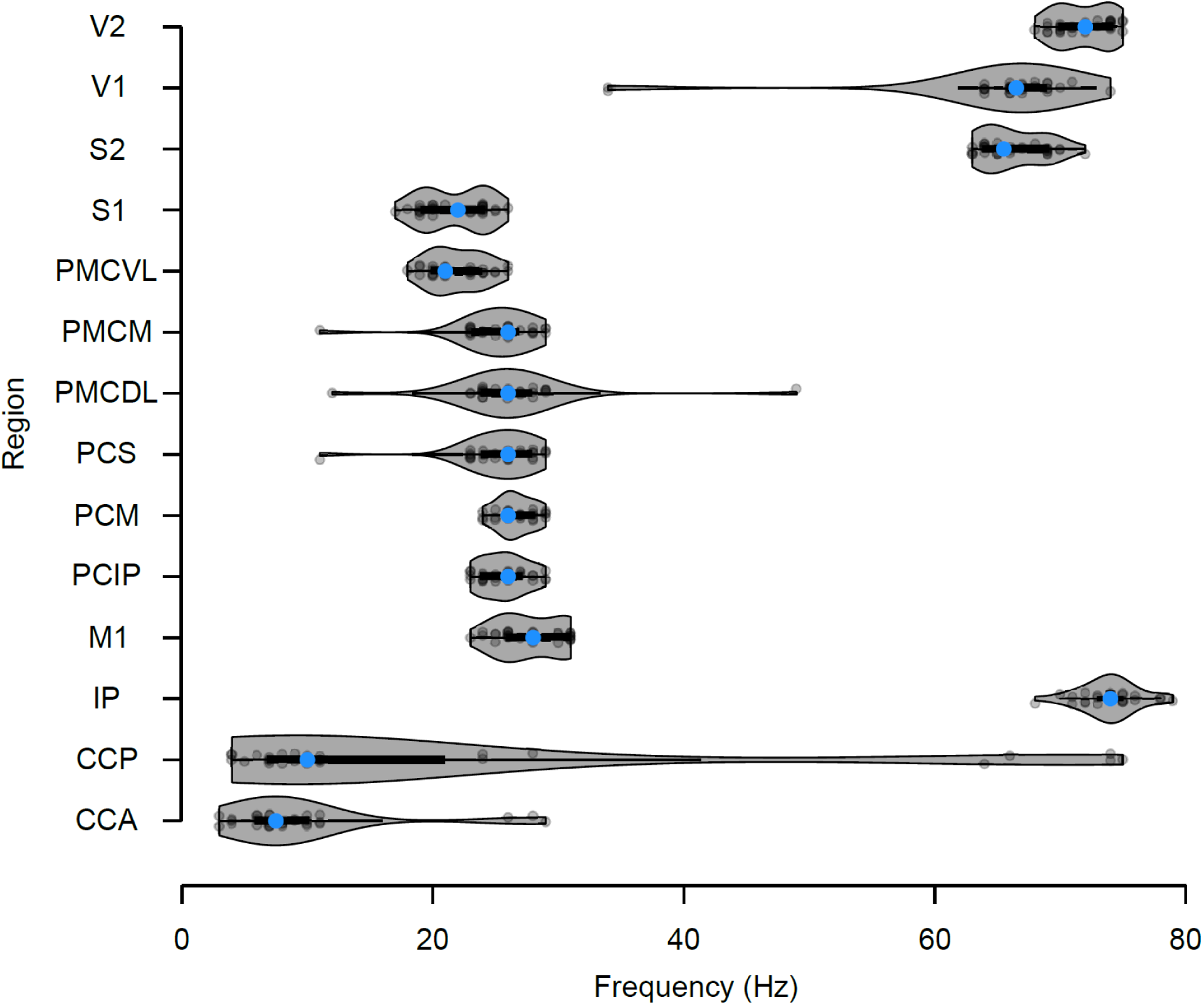
Most model brain region’s continued to produce the experimentally observed frequency band regardless of the *v̇*, *u̇* pair used to generate the region’s inherent activity. Violin plots of each brain region’s oscillation frequency in the complete visuomotor functional subnetwork for each of the 30 subnetwork instantiations. In all cases the oscillation frequencies varied with *v̇*, *u̇* selection, but for most cases, each region’s frequency remained in the experimentally-observed frequency band. For V1, PMCM, PMCDL, and PCS, one or two cases had frequencies outside the experimentally-observed band. For CCP and CCA, multiple anomalous frequencies occurred.

For most brain regions (V2, S2, S1, PMCVL, PCM, PCIP, M1, and IP), region oscillation frequency in the 30 visuomotor models, although showing some variability with *v̇*, *u̇* pair, was nonetheless always within the experimentally observed frequency band. For four of the regions (V1, PMCM, PMCDL, PCS) in one or two instantiations the region’s oscillation periods were outside the experimentally observed range. These anomalous frequencies were approximately half or twice the experimentally observed frequencies, suggesting that they were cases of “wrong” 2:1 or 1:2 coupling in the visuomotor model. For CCP (4 cases) and CCA (3 cases), alternatively, multiple instantiations had very anomalous frequencies. Nonetheless, these data show that in most instantiations (approximately 180 of 196 region activities, 92%), model brain region frequencies remained in the experimentally observed range regardless of the *v̇*, *u̇* pair used to produce the region’s inherent activity.

## 4 Discussion

We used here a novel modularity algorithm to identify a 14-brain-region functional visuomotor subnetwork from a primate whole-brain connectome. Local graph theory analyses of the subnetwork suggested that PCIP, V2 and PCS are centrally important regions for subnetwork activity. When populated by brain region models with appropriate inherent oscillatory activity and interconnected, the subnetwork produced activity similar to experimental data.

### 4.1 Important regions, some non-obvious, captured by modularity algorithm

In addition to capturing regions having obvious roles in visual and motor function (V1; V2; the premotor cortices PMCVL, PMCM, PMCDL; M1), the algorithm found several brain regions known from other work to be important in visuomotor function. In macaques, PCIP is integral in transforming sensory information into motor information for motor planning, particularly for arm and eye movements (Grefkes and Fink, 2005). The inner perisylvian region, which contains S2 and IP, acts as a sensorimotor integration area during hand manipulation. Some neurons from these areas are active before hand-target contact and others during finger exploration and precision grasping (Ishida et al., 2013). The cingulate cortex helps interpret outcome information of a present task to guide subsequent action selection. This region contains the anterior cingulate cortex (CCA), comprised of three motor areas, at least one of which receives inputs from adjacent motor areas and the prefrontal cortex, and all of which directly connect to M1, making it suitable for motor action selection when confronted with stimulus-task-reward behaviors (Walton and Mars, 2007) such as reach and grasp.

### 4.2 Regions structurally positioned to play an important role in visuomotor function

PCIP was identified in this work as the primary hub of the structural visuomotor subnetwork and scored lowest on the local resilience measure. It was thus structurally positioned to shape information flow throughout the network. PCIP is part of the dorsal stream of visual information and is the interface between the visual system identifying where objects to manipulate are and the motor system which uses the visual information to plan and execute the manipulation. Nakamura et al. (2001) showed in monkeys that feed-forward network connections between the association visual area, the lateral intraparietal area (lateral part of the PCIP), and the anterior intraparietal area (anterior part of the PCIP), where the latter projects to premotor areas, are important for transforming 3D visual information into proper hand movements. These data suggest that PCIP is the interface between visual and sensory systems and likely important for visuomotor integration and transformation.

Humans with cortical lesions in the anterior lateral bank of the PCIP have deficits in fine finger movements and, to a lesser degree, reaching movements (Binkofski et al., 1998). Tunik et al. (2005) continued this work, showing that transcranial magnetic stimulation creating virtual lesions of the anterior portion of the PCIP disrupted visually guided prehensile movement. Thus, consistent with its resilience network measure in our visuomotor functional subnetwork, altering PCIP activity compromises the brain’s ability to produce proper visuomotor function.

V2 was identified as another secondary hub and as the region with the third lowest local resilience score, making this region central to overall network function. This is consistent with histological studies of V2 projections in macaques, which show V2 reciprocal projections back to V1 and feed-forward projections to V3, V4, the middle temporal visual area, and medial superior temporal, parieto-occipital, and ventral intraparietal areas. V2 thus is widely connected to regions in both visual information streams (Gattass et al., 1997). V2 is also part of the prestriate cortex, the downstream brain area to which the striate cortex, containing V1, projects.

The local network analysis identified PCS as a secondary hub of the visuomotor subnetwork and the fourth lowest local resilience score. This region receives afferent projections from regions in the dorsal visual stream (e.g., the medial bank of the PCIP), parietal areas (e.g., different subregions of the PCS) and frontal motor association, premotor, and M1 areas, as well as from insular and cingulate areas, suggesting a role in goal-directed motor movement like arm movement and motor coordination (Bakola et al., 2010; Bakola et al., 2013). PCS activity changes with changing movement direction, dynamic position of the hand, hand movement preparation, and self-observation and monitoring of motion and/or position of the hand in the visual field (Battaglia-Mayer et al., 2001; Ferraina et al., 2001). PCS activity also encodes arm-movement direction and depth, with directional signals preceding arm movement (i.e., is responsible for proper motor planning) and depth-based signals being prominent during and after movement execution (De Vitis et al., 2019).

### 4.3 Limitations of current work

The primate whole-brain connectome used here contains only 76 brain regions. With increased time presumably more, perhaps many more, brain regions will be identified and connectomes made available. The algorithm described here is general and can function with essentially any brain region number. Having a connectome dataset with increased granularity would increase subnetwork resolution and allow finer local-graph-theory analysis. However, with respect to functional relevance (e.g., Figure 10), such increases are only valuable if an associated increase in LFP or similar measures of the activity of new regions are also obtained.

Another concern is the unit oscillator model used to model region activity. This approach assumes that the average activity of a brain region is the most important metric of the information the region is receiving, altering, and sending on. It is thus similar to arguing that metrics of total telephone calls between cities measure the nature of the information being transmitted by the speakers making the calls. This analogy makes this approach seem ridiculous, but it is important to note that individual neurons receive and transmit much less information than humans. It is thus possible that grouped activity of each brain region is a useful metric of information flow.

Even if so, another potential difficulty with the brain region models is the seemingly simple 2D formalism underlying the activity, which contains none of the multiple conductances, each with their own voltage and calcium sensitivities and time constants, present in real neurons. However, it is important to note that the 2D model’s simplicity is truly only seeming. Even the 2D version used here can produce complicated neuron responses such as post-inhibitory rebound. Relatively small additions endowing the model with multiple time constants would effectively reproduce the wide dynamic range present in real conductances.

A final concern is the simplicity of the inter-region synaptic connectivity, the *η* in Section **3.4.1 Linking brain region activities** in the Results, which does not incorporate the non-linear summation of simultaneous inhibitory and excitatory inputs (e.g., shunting of excitatory post-synaptic currents, different post-synaptic current equilibrium potentials). As with modeling brain region (unit) activity with a simple model, it is unclear how great this difficulty is. Post-synaptic potentials are slow and add according to the electrical distance from the synapse to the cell body. This temporal and spatial filtering makes it likely that some simplified function can represent reasonably well the synaptic input:output relationship.

In summary, although somewhat more complicated unit and synaptic connection functions should undoubtedly be used in future work, the greatest concern is the assumption that brain regions can be modeled as single units. However, until neurobiological techniques reach a point at which the activity of most or all neurons in brain regions can be individually monitored, the unit assumption is the best that can be made.

### 4.4 Extensions of present approach and general applicability

We modeled subnetwork activity only at steady-state. Real subnetwork function occurs over time, with the input activating a spreading set of regions until the motor response finally occurs. The duration between activating sensory input and motor response is short and it is possible, perhaps likely, that at no point in the process does any part of the subnetwork achieve steady-state. The brain region models have complicated transient routes to steady-state activity and, with the parameters used here, approach steady-state relatively slowly compared to physiological input:output durations. Moreover, even when quiescent (the hyperpolarized, single state condition in Figure 9A), the brain region model can respond to synaptic input with highly non-linear responses (e.g., responding to a transient inhibition with a subsequent large depolarization; in neurobiology jargon, a post-inhibitory rebound). As such, a next step in this work is to populate the subnetwork with some or all non-oscillatory brain region models to test whether region and synaptic parameters can be found in which transient activation of the input region can reproduce the experimental recordings of subnetwork activity.

With respect to application to other behaviors, the algorithm presented here should be able to identify a relevant functional subnetwork for any behavior with clear input and output brain regions and hence most to all motor acts triggered by sensory input. To our knowledge, the algorithm described here is novel, and may thus be a useful addition to the toolkits available for identifying functional brain region subnetworks that create classes of behaviors.

## 6 Data availability statement

The data analyzed in this study is subject to the following licenses/restrictions: US NASA NNX14AE73G. Requests to access these datasets should be directed to RE

## 7 Author contributions

Conceptualization, RE, MM, and SLH; methodology, RE; formal analysis, RE; investigation, RE; resources, MM, SAG, RKP, and SLH; data curation, RE and SLH; writing—original draft, RE; writing—review & editing, all authors; visualization, RE and SLH; supervision, MM, SAG, RKP, and SLH; project administration, MM, SAG, RKP, and SLH; funding acquisition, MM (NASA) and SLH (Ohio University).

## 9 Acknowledgements

The authors thank M. Day, P. Jung, A. Niemann, and T. Young for insightful comments on the research and presentation.

